# ER-tethering directs TREX1 penetration of a BAF-dependent barrier at micronuclei

**DOI:** 10.1101/2025.04.14.648782

**Authors:** Yanyang Chen, Eleonore Toufektchan, Xiaohan Luan, James Hickling, Roshan Xavier Norman, Paolo Cifani, Alex Kentsis, Wen Zhou, John Maciejowski

**Affiliations:** Molecular Biology Program, Sloan Kettering Institute, Memorial Sloan Kettering Cancer Center, New York, NY, 10065, USA; Southern University of Science and Technology, School of Life Sciences, Department of Immunology and Microbiology, Shenzhen, China; Molecular Pharmacology Program, Sloan Kettering Institute, Memorial Sloan Kettering Cancer Center, New York, NY, 10065, USA

**Author notes:** Corresponding authors: John Maciejowski, PhD Molecular Biology Program Sloan Kettering Institute Memorial Sloan Kettering Cancer Center New York, NY, 10065, USA.

**Keywords:** TREX1, cGAS, STING, micronuclei, chromothripsis, chromosomal instability

## Abstract

Micronuclei are membrane-encapsulated nuclear aberrations that form following chromosome segregation errors. Micronuclear membrane collapse permits access of the pattern recognition receptor cGAS and its antagonist, the TREX1 exonuclease. TREX1 carboxy-terminal domain mediated endoplasmic reticulum tethering association is essential for invasion into ruptured micronuclei, however the mechanisms underlying this dependency are unknown. Here, we identify barrier-to-autointegration nuclear assembly factor 1 (BAF) as a key regulator of TREX1 activity at micronuclei. BAF accumulates on micronuclei following membrane collapse and augments TREX1 recruitment in a manner that depends on BAF interactions with membrane-associated LEM-domain proteins. Despite delayed entry, TREX1 exhibits enhanced micronuclear DNA degradation and independence from ER-tethering in BAF-deficient cells. In accordance, recombinant BAF protein inhibits TREX1-mediated DNA degradation *in vitro* in a manner that depends on BAF DNA-binding. BAF similarly outcompetes cGAS for micronuclear DNA interaction and reduces cGAS activation at micronuclei. These findings reveal, a BAF-dependent protective barrier to diffusive entry of DNA binding proteins at ruptured micronuclei, explaining the requirement of TREX1 ER-tethering for micronuclear localization and suppression of productive cGAS DNA substrate interactions that activate innate immune responses in chromosomally unstable cells.

## INTRODUCTION

Chromosomal instability (CIN) is characterized by high rates of chromosome mis-segregation during cell division^1^ and resultant micronuclei formation when a chromosome or chromosome fragment fails to join the primary nucleus^2^. Micronuclei frequently lose compartmentalization due to irreversible membrane collapse^3,4^. Micronuclear membrane rupture can elicit an immune response by enabling the cytosolic DNA sensor cGAS to productively engage exposed, genomic, double-stranded DNA^5–9^. Although relationships to tumorigenesis are multi-faceted, sensing of cancer-intrinsic cytosolic DNA by the cGAS-STING signaling pathway can slow tumor growth by provoking antitumor immune responses^5,10–12^.

Chromosomally unstable cancer cells depend on adaptive mechanisms to evade cGAS-STING-dependent immune responses triggered by high levels of cytosolic DNA^13–17^. Upregulation of the TREX1 3′→5′ DNA exonuclease, a key cGAS-STING antagonist, shields chromosomally unstable tumors from immune surveillance by dampening Type I interferon (IFN) production^11,18–20^. Recognition of the critical role of TREX1 as an intratumoral checkpoint against innate immune activation has spurred therapeutic efforts to target TREX1 enzymatic activity^21,22^.

The critical nature of TREX1 function is demonstrated by its role in a spectrum of human diseases including the pediatric lupus like Aicardi-Goutières syndrome (AGS), systemic lupus erythematosus, and retinal vasculopathy with cerebral leukodystrophy^23–26^. Biallelic mutations that compromise TREX1 nucleolytic activity are predominantly associated with AGS, an immune disease characterized by neurological dysfunction and high systemic levels of type I IFN^23,27^. *Trex1* KO and TREX1-nuclease deficient mice recapitulate the hallmarks of AGS^28–32^. The AGS-like phenotypes of *Trex1*-deficient animals are reversed by deletion of cytosolic DNA sensing components, including *cGas*, *Sting1*, and *Ifnar1,* thus implicating deficient cytosolic DNA degradation and chronic cGAS activation in the etiology of AGS^28,33,34^. In contrast, autosomal dominant mutations that truncate the C-terminus of TREX1 disrupt its association with the ER, while preserving nucleolytic activity, are associated with the small vessel disease retinal vasculopathy with cerebral leukoencephalopathy^35^.

Here, we report that barrier-to-autointegration nuclear assembly factor 1 (BAF) plays critical roles in regulating TREX1 activity at ruptured micronuclei. BAF engages micronuclei shortly after membrane collapse in a manner that depends on its ability to bind DNA. Working together with partner LEM domain proteins ANKLE2 and LEMD2, BAF promotes entry of TREX1-ER to ruptured micronuclei. Despite delayed recruitment to micronuclei, TREX1 catalyzed higher levels of micronuclear DNA resection in a manner independent of ER-tethering in BAF-depleted cells. Our results indicate that ER-tethering enables TREX1 to overcome BAF-mediated protection of micronuclear DNA. This active form of transport to micronuclei is necessary to limit cGAS activation of innate immune responses in chromosomally unstable cells.

## RESULTS

### Proteomic analysis of ruptured micronuclei identifies enrichment of BAF and LEM domain proteins

To investigate mechanisms that could potentially regulate TREX1 activity at micronuclei, we characterized the proteomes of intact and ruptured micronuclei to identify differences in protein composition (Figure 1A). TREX1 localizes to micronuclei shortly following membrane collapse^8,36^. We therefore used a previously developed strategy to isolate micronuclei based on their association with cGAS, a well-established marker of ruptured micronuclei^5,6,8,37^. In brief, HEK293T or HeLa cells were labeled with H2B-mCherry and GFP-cGAS to mark chromatin and ruptured micronuclei respectively (Figure 1A). Lysates were then fractionated by differential centrifugation and micronuclei-containing fractions were further processed by flow cytometry to isolate micronuclei based on GFP-cGAS labeling intensity (Figure 1B). As previously reported^8^, all purified micronuclei exhibited a similar distribution of area, while the mean GFP-cGAS signal intensity differed more than 2-fold across intact and ruptured micronuclei populations (Figures S1A-C). Purified nuclei and micronuclei were then analyzed using label-free quantitative high-resolution mass spectrometry proteomics. As expected, based on its central role in our experimental workflow, comparative proteomic analysis confirmed a strong enrichment of cGAS in ruptured micronuclei relative to intact micronuclei or primary nuclei purified from HeLa cells (Figures 1C and 1D; Table S1). Consistent with our prior results^8^, (3xFLAG)-TREX1 was enriched in micronuclei relative to primary nuclei purified from HEK293T cells (Figures S1D). RPA1, a component of the RPA complex that associates with micronuclear ssDNA following TREX1-mediated resection^8^, showed a similar enrichment at ruptured micronuclei relative to intact micronuclei (Table S1).

**Figure 1.**
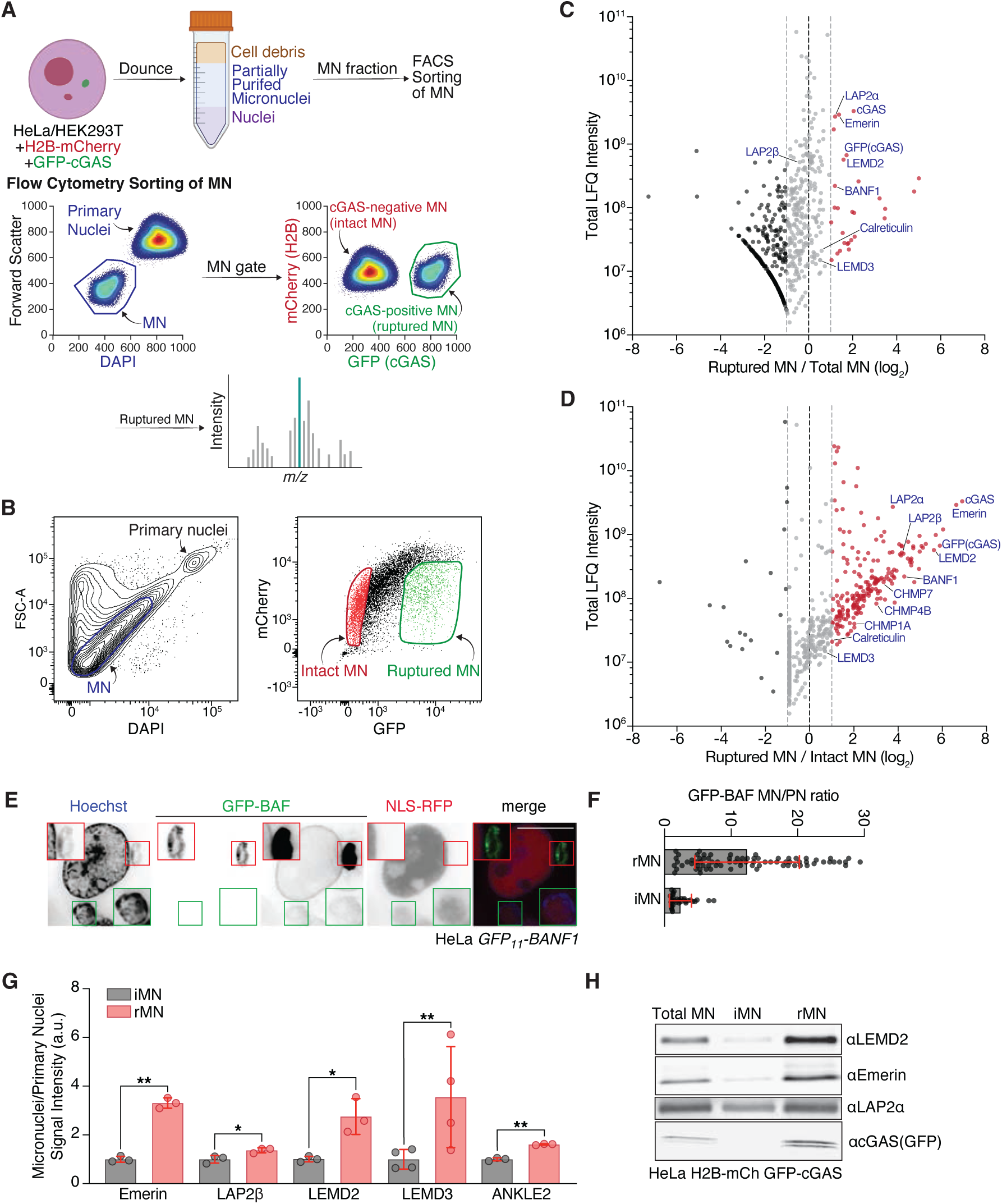
Proteomics identifies enrichment of BAF and LEM domain proteins in ruptured micronuclei. (A) Schematics of micronuclei purification workflow. (B) Flow profiles of partially purified micronuclei from HeLa H2B-mCherry GFP-cGAS cells. (C) Volcano plots showing relative protein abundances in ruptured versus total micronuclei in HeLa H2B-mCherry GFP-cGAS cells. Displayed proteins were identified at FDR<0.01. Relative abundance is expressed as log₂-transformed ratio of the total area-under-the-curve for peptides assigned to each protein in different conditions. Total LFQ intensities (Label-Free Quantification, au) assigned to each protein is used as surrogate for signal-to-noise and accuracy of quantification. GFP-cGAS is used as a marker for ruptured micronuclei. Two independent experiments were performed. (D) Volcano plots showing relative protein abundances in ruptured versus intact micronuclei, as in (C). (E) Live cell imaging of endogenously tagged HeLa GFP_11_-BAF GFP_1-10_ cells. Scale bar, 10 µm. (F) Quantification of GFP-BAF micronuclei over primary nuclei signal intensity ratio as shown in (E). Mean ± s.d. of individual cells from *n* = 1 experiment are shown. (G) Quantification of LEM domain proteins micronuclei over primary nuclei signal intensity ratio as shown in Figure S1E. Mean ± s.d. of *n* = 3 experiments are shown with greater than 100 total micronuclei quantified per experiment. *P* values were calculated by Student’s *t*-test (**P < 0.01, *P < 0.1) (H) Immunoblotting for the indicated proteins in MN sorted from HeLa H2B-mCherry GFP-cGAS cells. See also Figure S1.

Further inspection of the mass spectrometry data identified additional proteins previously reported to accumulate at ruptured micronuclei. These include components of the ESCRT-III complex, such as CHMP1A, CHMP4B, and CHMP7 (Figures 1D; Table S1), which aberrantly accumulate at micronuclei where they drive membrane deformation and catastrophic membrane collapse^38,39^. Consistent with prior observations of ER tubule invasion into micronuclear chromatin following membrane collapse, proteomic analyses confirmed an enrichment of ER-resident proteins, such as Sec61, RPN1, and calreticulin in ruptured micronuclei relative to intact micronuclei (Figure 1D; Table S1). Prior proteomic profiling of purified micronuclei demonstrated that spliceosome machinery was depleted from micronuclei induced by genotoxic stress, a defect associated with micronuclear DNA damage^40^. These prior observations were also reflected in our proteomic data (Table S1), which showed a depletion of spliceosome components, including Prp3, SCAF1, and SNRPB2, in ruptured micronuclei relative to intact micronuclei. Together, these data confirm the accuracy of MS-based proteomic profiling to characterize the proteomes of purified micronuclei.

Barrier-to-autointegration nuclear assembly factor 1 (BAF) was among the proteins showing the strongest enrichment on ruptured micronuclei relative to intact micronuclei (19-fold) (Figure 1D; Table S1). BAF is a small, DNA-binding protein that rapidly localizes to sites of nuclear envelope rupture, including ruptured micronuclei^4,41–43^. In addition to BAF, proteomic profiling revealed an enrichment of LAP2-emerin-MAN1 (LEM) domain proteins, a family of nuclear lamina proteins characterized by a ∼40 amino acid LEM domain that binds to BAF^44^, at ruptured micronuclei relative to intact micronuclei (Figure 1D; Table S1). Altogether, ruptured micronuclei exhibited an enrichment of four components of the seven-member LEM domain family, including emerin, LAP2α, LAP2β, and LEMD2.

To further investigate BAF localization we integrated a fragment of GFP consisting of the eleventh β-strand of super-folder GFP (GFP_11_) into the endogenous *BANF1* locus (Figures S1E and S1F). Complementation of GFP by transgenic overexpression of the remaining GFP_1-10_ fragment restored fluorescence thus allowing for visualization of BAF subcellular localization^45^. Consistent with our proteomic data and published findings^4,43^, live-cell imaging confirmed an approximately 10-fold enrichment of BAF on ruptured micronuclei relative to intact micronuclei (Figures 1E and 1F). Immunofluorescence analysis of LEM domain protein localization revealed similar 3-4-fold increases of emerin, LAP2β, and LEMD2 signal intensity at ruptured micronuclei (Figure 1G; Figure S1G). Although not detected in our proteomic analyses, anti-LEMD3 and anti-ANKLE2 immunofluorescence revealed similar increases in localization to ruptured micronuclei (Figure S1G). Immunoblotting analysis of purified micronuclei further confirmed enrichment of emerin, LAP2α, and LEMD2 at ruptured micronuclei (Figure 1H). ANKLE1, a LEM domain endonuclease proposed to process chromatin bridges^8,46^, could not be detected at micronuclei, potentially due to low expression levels in HeLa cells (Figure S1H). Thus, together with its partner LEM domain proteins, BAF exhibits a strong enrichment at ruptured micronuclei.

### BAF promotes TREX1 recruitment to ruptured micronuclei

We next sought to understand how proteins enriched at ruptured micronuclei affected TREX1 recruitment to micronuclei following micronuclear membrane collapse. To study TREX1 localization dynamics we integrated a GFP_11_ tag into the endogenous *TREX1* locus (Figures 2A and 2B; Figures S2A and S2B). As above, this approach allowed us to follow TREX1 subcellular localization through complementation of GFP by transgenic overexpression of the remaining GFP_1-10_ fragment. As predicted by prior analyses of GFP-TREX1 overexpression and anti-TREX1 immunofluorescence^8,28,36,47^, live-cell imaging of GFP_11_-TREX1 showed strong colocalization with the ER and enrichment in >60% of ruptured micronuclei (Figures 2C and 2D; Figure S2C). Importantly, ELISA measurements of cGAMP production showed that GFP_11_-TREX1 was indistinguishable from wild-type TREX1 in terms of cGAS regulation, indicating that integration of the GFP_11_-tag did not disrupt normal TREX1 function (Figure S2D).

**Figure 2.**
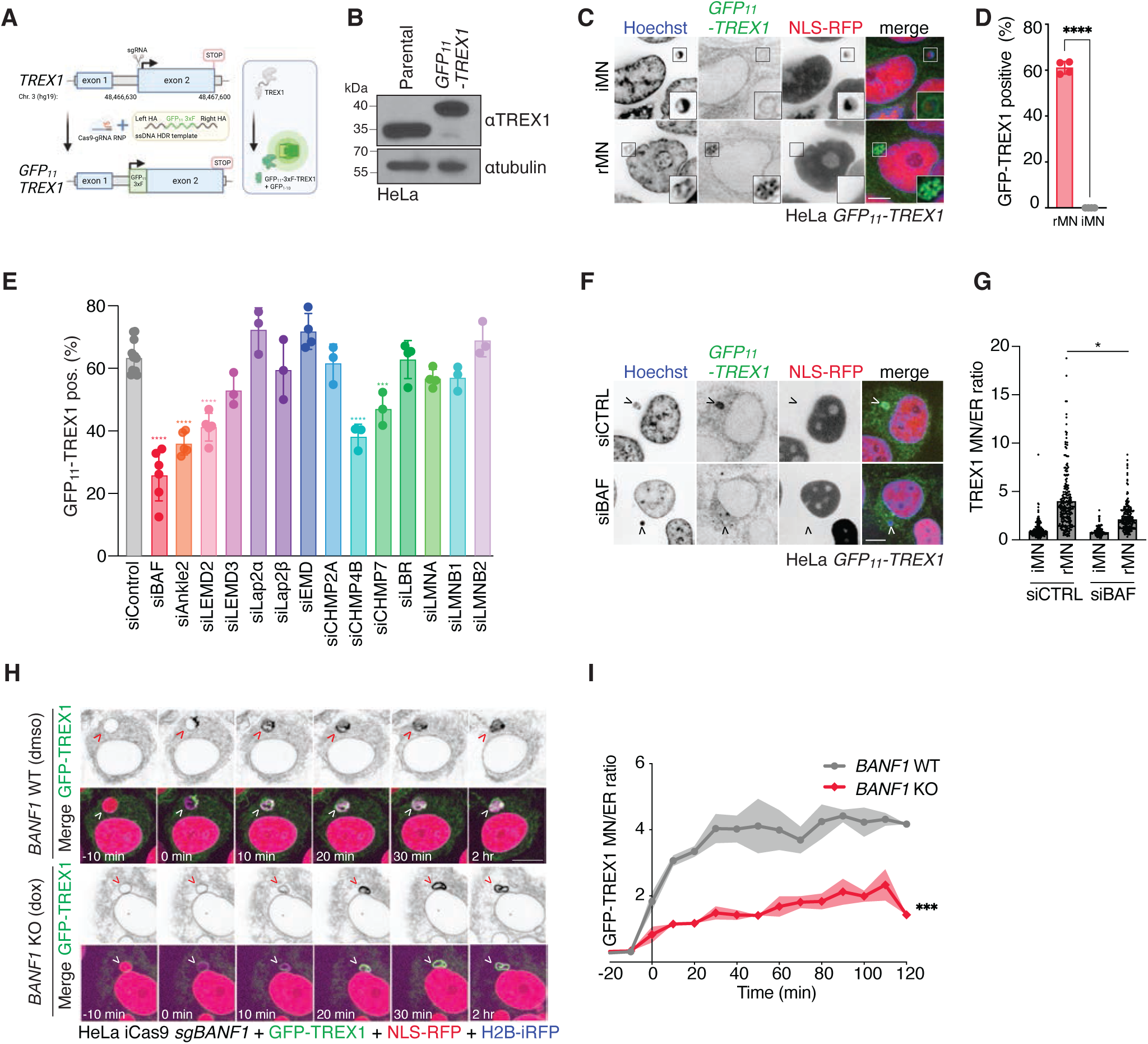
BAF promotes TREX1 recruitment to ruptured micronuclei. (A) Schematics of targeting GFP_11_ to TREX1 N terminus in HeLa GFP_1-10_ cells. (B) Immunoblotting of HeLa GFP_1-10_ GFP_11_-TREX1 with the indicated antibodies. (C) Live-cell imaging of HeLa GFP_1-10_ GFP_11_-TREX1 along with NLS-RFP. DNA was stained with Hoechst. Scale bar, 5 µm. (D) Quantification of the percentage of GFP-TREX1 positive micronuclei as shown in (C). Mean ± s.d. of *n* = 4 independent experiments with greater than 100 total micronuclei quantified per experiment. *P* value was calculated by Student’s *t*-test (****P < 0.0001). (E) siRNA screen quantification of percentage of GFP-TREX1 positive micronuclei to identify factors required for TREX1 recruitment to ruptured MN. Mean ± s.d.; ****P < 0.0001, ***P < 0.001, ordinary one-way ANOVA with Dunnett’s multiple comparisons test; ≥3 independent experiments for each siRNA treatment. From left to right: Total MN analyzed: *n* = 1589, 567, 494, 612, 342, 232, 295, 273, 278, 386, 358, 694, 465, 599, 387; Ruptured MN analyzed: *n* = 967, 441, 385, 453, 201, 169, 216, 183, 170, 268, 247, 360, 287, 298, 229. (F) Live-cell imaging of HeLa GFP_1-10_ GFP_11_-TREX1 NLS-RFP with BAF depletion by siRNA. Scale bar, 10 µm. (G) Quantification of the normalized fluorescence signal intensities of GFP-TREX1 MN/ER ratio in HeLa stably expressing GFP-TREX1 with BAF depletion by siRNA. Mean ± s.d. of *n* = 3 independent experiments are shown with greater than 100 total micronuclei quantified per experiment. *P* value was calculated by Student’s *t*-test (*P < 0.1). (H) Live-cell time-lapse imaging of HeLa inducible BAF KO stably expressing GFP-TREX1 after 72 hours treatment of dmso (*BANF1* WT) or doxycycline (*BANF1* KO), captured at 10-minute intervals. Time 0 min marks the micronuclear envelope rupture, indicated by the loss of NLS-RFP signal. Scale bar, 10 µm. (I) Quantification of the normalized fluorescence signal intensities of GFP-TREX1 MN/ER ratio as shown in (H). Mean ± s.e.m. ***P<0.001. Wilcoxon matched-pairs signed rank test; 3 independent experiments. *BANF1* wt n = 23 ruptured MN; *BANF1* KO *n* = 26 ruptured MN. See also Figure S2.

Next, we performed a targeted siRNA screen to investigate how depletion of micronuclear proteins affects GFP_11_-TREX1 recruitment to ruptured micronuclei. The screen was composed of factors showing an enrichment at ruptured micronuclei including BAF, LEM domain proteins (Ankle2, LEMD2, LEMD3, LAP2α, LAP2β, Emerin) and ESCRTs (CHMP2A, CHMP4B, CHMP7) (Figure 2E; Figure S2E). Additional nuclear membrane proteins not showing an enrichment at ruptured micronuclei, such as LBR, Lamin A, Lamin B1, and Lamin B2, were included as controls. Following siRNA transfection, micronuclei were induced by nocodazole washout and GFP_11_-TREX1 subcellular localization was tracked by live-cell imaging. Micronuclear envelope integrity was monitored by loss of NLS-RFP signal^3^. Once again, >60% of ruptured micronuclei were marked as GFP_11_-TREX1 positive indicating accumulation of GFP_11_-TREX1 at ruptured micronuclei and its apparent intercalation throughout micronuclear chromatin (Figure 2E). Consistent with a prior report^39^, depletion of the ESCRT-III subunits CHMP4B and CHMP7, decreased GFP_11_-TREX1 localization to <40% of ruptured micronuclei likely reflecting membrane deformations caused by unrestrained ESCRT-III activity (Figure 2E; Figure S2F).

However, the strongest decreases in GFP_11_-TREX1 localization to ruptured micronuclei were observed following depletion of BAF and LEM domain proteins Ankle2 and LEMD2 (Figure 2E; Figure S2G). Quantification of GFP_11_-TREX1 signal intensity confirmed a >2-fold decrease in the percentage of GFP_11_-TREX1 positive ruptured micronuclei following BAF depletion (Figures 2E and 2F). Of note depletion of BAF, Ankle2, and LEMD2 led to modest increases in micronuclear envelope rupturing rates likely due to roles in preserving nuclear membrane stability (Figure S2H)^4^. Importantly, TREX1 protein levels were unaffected by siBAF transfection indicating that diminished TREX1 localization to ruptured micronuclei reflects a defect in recruitment as opposed to a reduction in TREX1 protein stability (Figure S2I).

To further study the impact of BAF depletion on TREX1 localization to ruptured micronuclei we used dox-inducible Cas9 expression to conditionally delete *BANF1* in HeLa cells (Figures S2J and S2K). Consistent with our siRNA screening data, time-lapse, live-cell imaging revealed that *BANF1* deletion delayed, but did not completely disrupt, GFP-TREX1 localization to ruptured micronuclei (Figures 2H and 2I; Figure S2L; Videos S1 and S2). Taken together, these data indicate that, along with its partner LEM domain proteins, BAF plays an essential role in the timely recruitment of TREX1 to ruptured micronuclei.

### BAF-LEM domain interactions are necessary for TREX1-ER recruitment

BAF recruits ER membranes to chromosomes during reassembly of the nuclear lamina at mitotic exit and to rupture sites at the primary nucleus^41,43,48,49^. These activities aid in nuclear envelope reassembly and repair and the restoration of compartmentalization. We therefore used time-lapse, live-cell imaging to monitor how BAF depletion affects ER membrane recruitment to ruptured micronuclei. As previously reported^3,8,39^, ER tracker-labeled membranes invaded micronuclei immediately following rupture, albeit without the strong enrichment exhibited by TREX1 relative to neighboring ER (Figures 3A and 3B; Figure 2H). In contrast, ER membranes remained excluded from micronuclei for the duration of imaging, up to 2 hrs following rupture, in BAF-depleted cells (Figures 3A and 3B). Thus, similar to its functions at ruptures of the primary nucleus, BAF promotes ER membrane recruitment to ruptured micronuclei.

**Figure 3.**
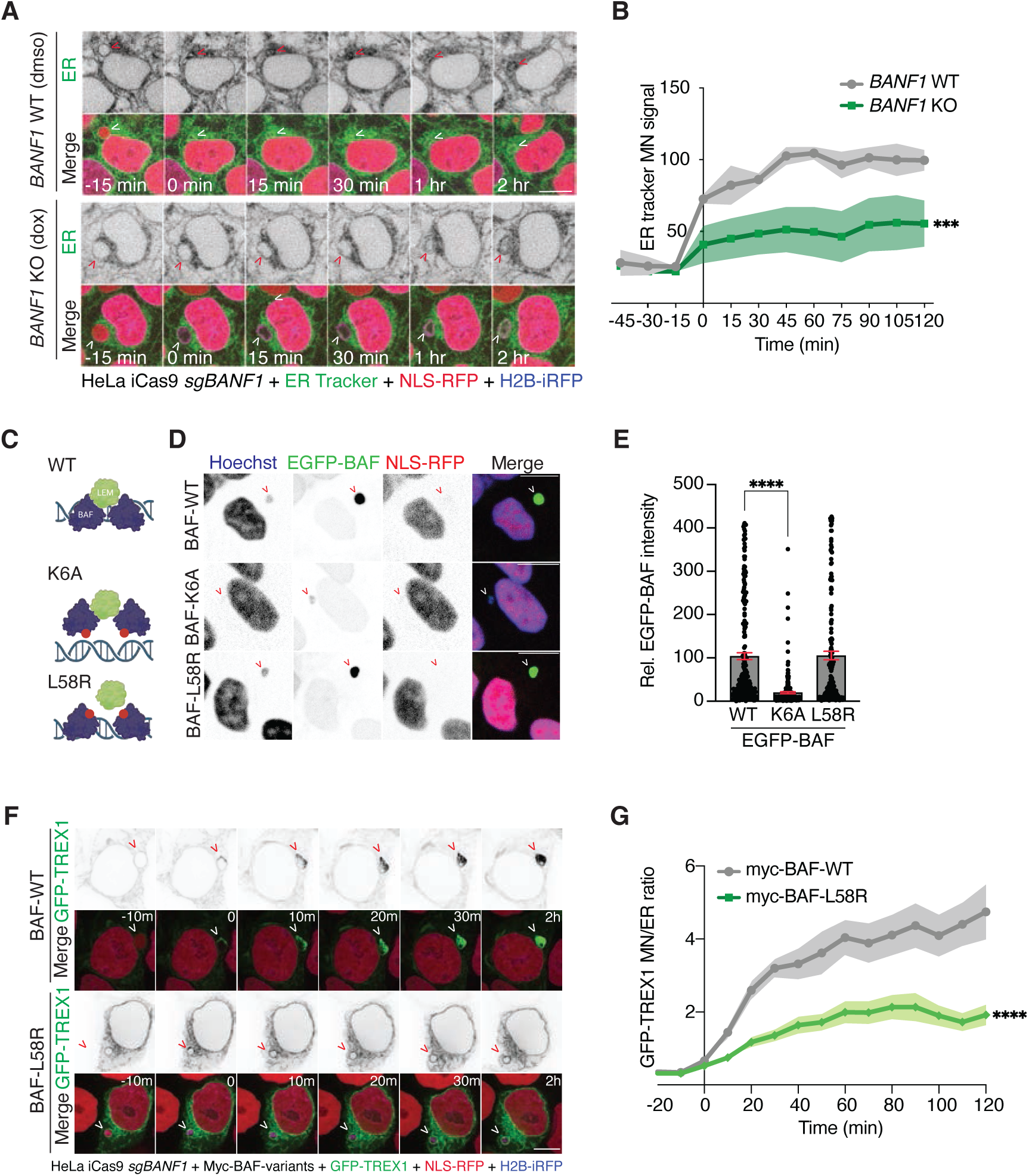
BAF-LEM domain interactions are necessary for TREX1-ER recruitment. (A) Live-cell time-lapse imaging of HeLa inducible *BANF1* KO after 72 hours treatment of dmso (*BANF1* WT) or doxycycline (*BANF1* KO) and stained with ER tracker Green, captured at 10-minute intervals. Arrow indicated a micronucleus going through rupture. Time 0 min marks the micronuclear envelope rupture, indicated by the loss of NLS-RFP signal. Scale bar, 10 µm. (B) Quantification of the normalized fluorescence signal intensities of ER tracker as shown in (A). Mean ± s.e.m. ***P<0.001. Wilcoxon matched-pairs signed rank test; 3 independent experiments. *BANF1* wt n = 34 ruptured MN; *BANF1* KO *n* = 51 ruptured MN. (C) Schematics of BAF separation of function mutants. (D) Immunofluorescence of HeLa parental cells stably expressing EGFP-BAF-variants and NLS-RFP. Arrows marked ruptured micronuclei indicated by the loss of NLS-RFP signal. Scale bar, 10 µm.(E) Quantification of normalized EGFP-BAF intensity at ruptured micronuclei as shown in (D). Mean and SEM of n = 3 independent experiments with greater than 100 total micronuclei quantified per experiment. Ordinary one-way ANOVA with Dunnett’s multiple comparisons test (****P < 0.0001). (F) Live-cell time-lapse imaging of HeLa inducible *BANF1* KO stably expressing GFP-TREX1 rescued by Myc-BAF-WT or L58R after 72 hours treatment of doxycycline (*BANF1* KO), captured at 10-minute intervals. Time 0 min marks the micronuclear envelope rupture, indicated by the loss of NLS-RFP signal. Scale bar, 10 µm. Scale bar, 10 µm. (G) Quantification of the normalized fluorescence signal intensities of GFP-TREX1 MN/ER ratio as shown in (F). Mean ± s.e.m. ****P<0.0001. Wilcoxon matched-pairs signed rank test; *n* = 3 independent experiments. BAF WT *n* = 22 ruptured MN; BAF L58R *n* = 29 ruptured MN. See also Figure S3.

BAF binds to DNA directly through a helix-hairpin-helix DNA-binding domain that forms stable complexes on DNA, with estimated dissociation constants in the low femtomolar range^50^. To test whether DNA binding is necessary for BAF localization to ruptured micronuclei, we performed immunofluorescence to monitor the subcellular localization of a BAF K6A mutant with impaired DNA binding (Figure 3C)^51^. As expected, BAF-K6A mutants displayed severely reduced localization to ruptured micronuclei relative to wild-type controls (Figures 3D and 3E; Figure S3A). In contrast, BAF-L58R mutants, which are defective for LEM-domain binding^52^, accumulated normally at ruptured micronuclei. Thus, the ability of BAF to bind DNA is critical for its recruitment to ruptured micronuclei suggesting that BAF is attracted to micronuclei through cytosolic exposure of genomic DNA following micronuclear envelope rupturing.

We next sought to understand the mechanisms connecting BAF accumulation at ruptured micronuclei to TREX1-ER recruitment. BAF facilitates repair of primary nuclear envelope ruptures through recruitment of LEM-domain proteins to rupture sites^41,43^. Similar to these observations, BAF depletion led to reductions in the levels of LEM-domain proteins Ankle2 and LEMD2 at ruptured micronuclei (Figures S3B-E). To test if BAF LEM-domain binding is necessary for TREX1 recruitment to ruptured micronuclei, we performed time-lapse, live-cell imaging to monitor GFP-TREX1 subcellular localization in *BANF1* KO cells reconstituted with myc-BAF-wt or myc-BAF-L58R mutants (Figures 3F and 3G; Figure S3F). While myc-BAF-wt restored normal GFP-TREX1 accumulation to ruptured micronuclei, GFP-TREX1 remained excluded from ruptured micronuclei for the duration of imaging in cells reconstituted with myc-BAF-L58R mutants (Figures 3F and 3G). Thus, BAF promotes TREX1-ER recruitment to ruptured micronuclei in a manner that depends on its ability to interact with transmembrane LEM-domain proteins.

### Enhanced TREX1 DNA degradation and ER bypass in BAF-deficient cells

Following its accumulation at ruptured micronuclei, TREX1 accesses and resects micronuclear DNA resulting in ssDNA formation that can be monitored through RPA or native BrdU foci formation^8^. As previously reported, time-lapse, live-cell imaging of GFP-RPA revealed rapid foci formation in micronuclei as quickly as 10 minutes following micronuclear envelope rupturing (Figures 4A and 4B; Video S3). We reasoned that delayed recruitment of TREX1 to ruptured micronuclei in BAF-deficient cells may disrupt TREX1-dependent resection of micronuclear DNA thus resulting in delayed or decreased RPA foci formation. Indeed, as expected, GFP-RPA foci formation was significantly delayed in *BANF1* KO cells (Figures 4A and 4B; Video S4). However, longer time-lapse, live-cell imaging experiments identified strong increases in RPA foci formation at ruptured micronuclei suggesting that delays observed in *BANF1* KO cells may be temporary (Figures S4A; Videos S3 and S4).

**Figure 4.**
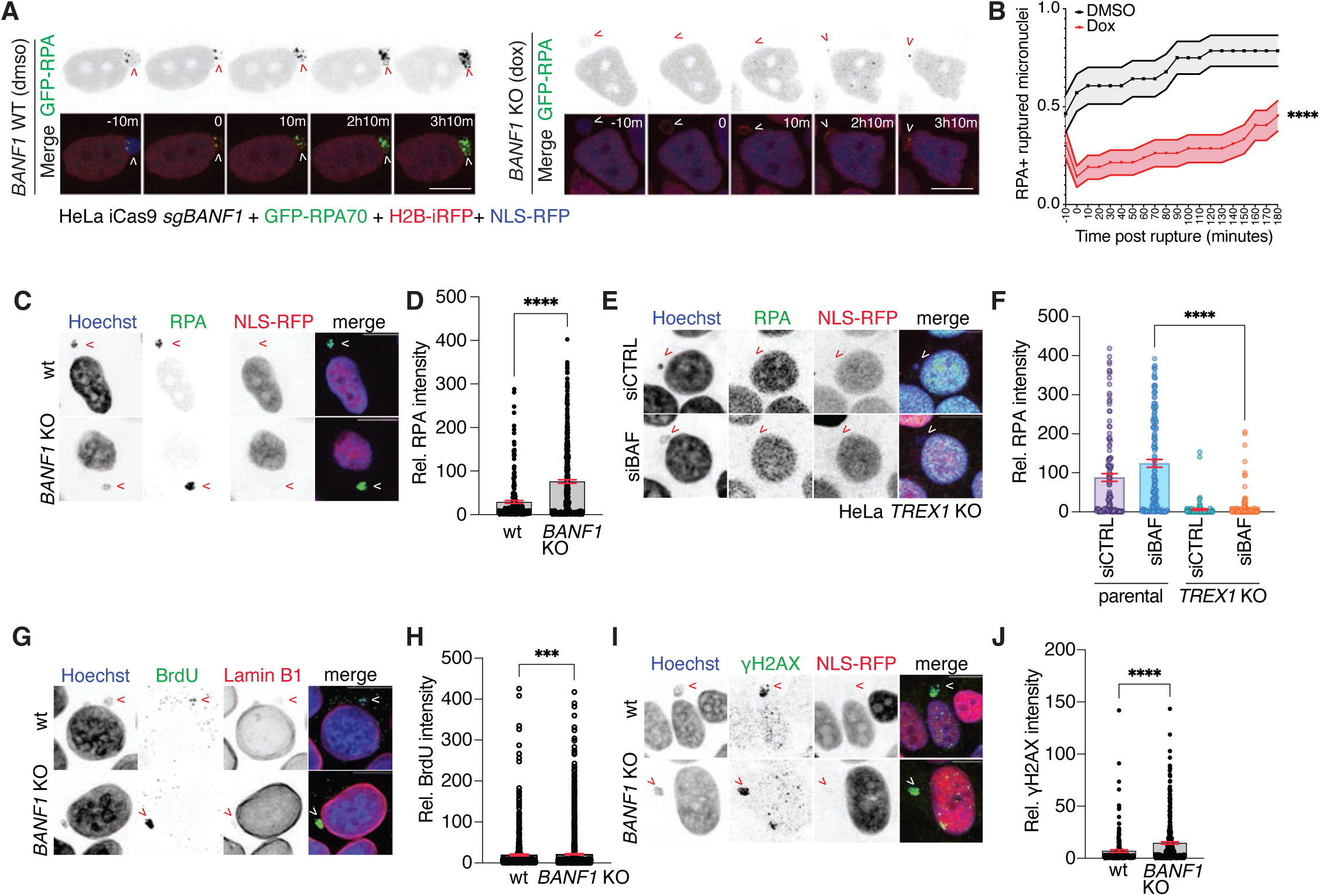
Enhanced TREX1 DNA degradation and ER bypass in BAF-deficient cells. (A) Live-cell time-lapse imaging of HeLa inducible BAF KO stably expressing GFP-RPA after 72 hours treatment of dmso (*BANF1* WT) or doxycycline (*BANF1* KO), captured at 10-minute intervals. Arrow indicated a micronucleus going through rupture. Time 0 min marks the micronuclear envelope rupture, indicated by the loss of NLS-RFP signal. Scale bar, 10 µm. (B) Quantification of the fraction of RPA + ruptured micronuclei as shown in (A). Mean ± s.e.m. ****P<0.0001. Wilcoxon matched-pairs signed rank test; 3 independent experiments. *BANF1* wt *n* = 28 ruptured MN; *BANF1* KO *n* = 42 ruptured MN. (C) Immunofluorescence of RPA in HeLa iCas9 sgBAF expressing NLS-RFP. Arrows marked ruptured micronuclei indicated by the loss of NLS-RFP signal. Scale bar, 10 µm. (D) Quantification of normalized RPA intensity in ruptured micronuclei as in (C). (E) Immunofluorescence of RPA in HeLa TREX1 KO treated with CTRL or BAF siRNA. Arrows marked ruptured micronuclei indicated by the loss of NLS-RFP signal. Scale bar, 10 µm. (F) Quantification of normalized RPA intensity in ruptured micronuclei as in (E). (G) Immunofluorescence of native BrdU in HeLa iCas9 sgBAF expressing NLS-RFP. Arrows marked ruptured micronuclei indicated by the loss of LaminB1 signal. Scale bar, 10 µm. (H) Quantification of normalized BrdU intensity in ruptured micronuclei as in (G). (I) Immunofluorescence of γH2AX in HeLa iCas9 sgBAF expressing NLS-RFP. Arrows marked ruptured micronuclei indicated by the loss of NLS-RFP signal. Scale bar, 10 µm (J) Quantification of normalized γH2AX intensity in ruptured micronuclei as in (I). *P* values in (D)(F)(H)(J) were calculated by Student’s *t*-test (***P < 0.001; ****P < 0.0001). Mean ± s.e.m. of *n* = 3 independent experiments with greater than 100 total micronuclei quantified per experiment. See also Figure S4.

To complement our time-lapse imaging analyses we next monitored ssDNA formation in ruptured micronuclei by quantifying RPA32 levels via immunofluorescence. This approach lacks the precise time resolution of time-lapse imaging but allows for examination of the long-term impacts of micronuclear envelope rupturing on micronuclear DNA resection^8^. Surprisingly, in contrast to our time-lapse imaging data, this analysis identified a nearly 3-fold increase in RPA32 mean signal intensity at ruptured micronuclei in *BANF1* KO cells (Figures 4C and 4D). Similar results were observed following BAF depletion by siRNA suggesting that increases in RPA32 signal intensity at ruptured micronuclei were not an artifact of any specific BAF depletion method (Figures S4B and S4C). Although normally dependent on TREX1^8^, we reasoned that BAF depletion may enable access to an additional nuclease that is normally excluded from ruptured micronuclei. However, *TREX1* deletion severely decreased RPA32 signal intensity at ruptured micronuclei regardless of BAF status (Figures 4E and 4F). Thus, the increased ssDNA levels at ruptured micronuclei observed following BAF depletion are a result of enhanced TREX1 micronuclear DNA resection.

To more directly assess micronuclear DNA resection we quantified 5-bromo-2′-deoxyuridine (BrdU) levels in MN under non-denaturing conditions, in which BrdU remains undetectable unless exposed by DNA resection^53^. This analysis showed that BrdU signal intensity was significantly increased following BAF depletion by CRISPR-Cas9 or siRNA relative to wild-type controls, thus further supporting a model of BAF-mediated protection against TREX1 DNA resection at ruptured micronuclei (Figures 4G and 4H; Figures S4D and S4E).

To further investigate how BAF impacts micronuclear DNA damage we performed immunofluorescence to assess γH2AX signal intensity at ruptured micronuclei. This analysis revealed 2.1-fold to 1.4-fold increases in γH2AX signal intensity in *BANF1* KO and BAF-depleted cells relative to wild-type controls (Figures 4I and 4J; Figures S4H and S4I). Although TREX1 appears to be primarily responsible for ssDNA increases observed following micronuclear envelope rupturing, it plays a more modest role in generating DNA damage as monitored by γH2AX foci, likely due to additional contributions from pathological base excision repair^8,54^. Nevertheless, *TREX1* deletion completely eliminated increases in γH2AX signal intensity observed following BAF depletion (Figures S4H and S4I). Altogether, these data indicate that BAF protects micronuclear DNA from ssDNA accumulation and DNA damage resulting from TREX1-mediated resection within ruptured micronuclei.

### BAF regulates TREX1 DNA degradation *in vitro*

We next moved to a purified system to understand how co-incubation with BAF affects TREX1-mediated DNA degradation *in vitro*. As previously reported, human TREX1 exhibited robust DNA exonuclease activity against a model dsDNA substrate (Figures 5A-D; Figures S5A). However, similar to our observations at ruptured micronuclei, TREX1 DNase activity was largely inhibited by high concentrations of BAF (>4 µM) (Figures 5A and 5B). This effect was specific to high concentrations of BAF, as BAF co-incubation enhanced TREX1 DNase activity at lower concentrations (1-2 µM) (Figures 5A-D). Similar results were observed using fish TREX1, which shares 26% homology to the human protein, indicating that BAF-mediated regulation of TREX1 DNase activity is conserved across evolution (Figures S5B). BAF K6A mutants failed to inhibit TREX1 activity, even at high concentrations, indicating that TREX1 inhibition by BAF depends on its DNA-binding ability (Figures 5B and 5D). Inhibitory effects of BAF on TREX1 activity appear to be unique, as an N-terminal fragment of cGAS and C-terminal fragment of OAS1 failed to inhibit TREX1 activity (Figures 5C and 5D; Figures S5C). Consistent with our prior observations^55^, cGAS N-terminal fragments formed higher-order complexes with DNA, likely reflecting a phase separation mechanism^56^, but failed to provide strong resistance against TREX1 nuclease activity at this concentration (10 µM). Thus, BAF inhibits TREX1 mediated DNA degradation *in vitro* in a mechanism that depends on its DNA-binding ability and relative concentration differences, explaining how TREX1 can still actively resect genomic DNA in BAF positive micronuclei.

**Figure 5.**
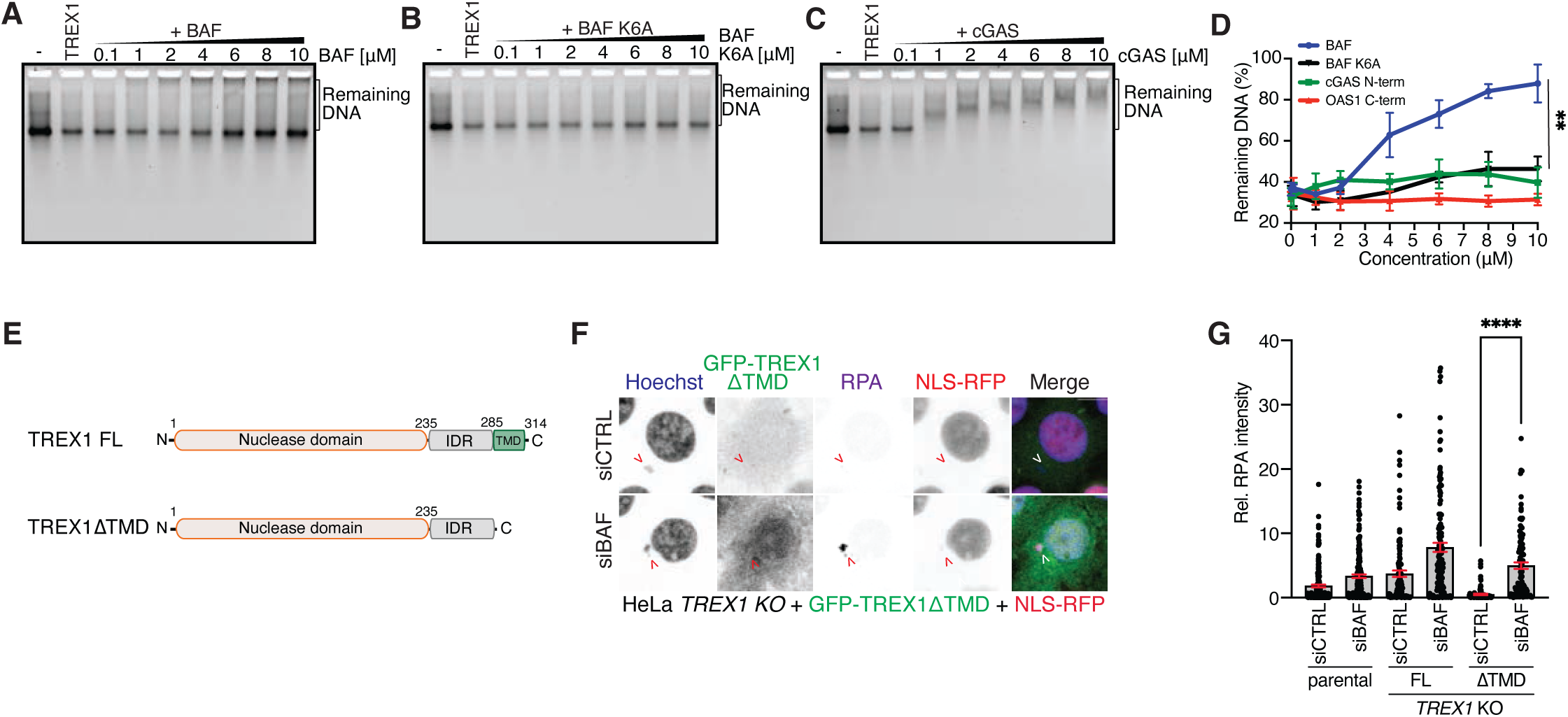
BAF regulates TREX1 DNA degradation *in vitro*. (A) Representative DNA gel from *in vitro* nuclease assay. A dsDNA substrate was co-incubated with a fixed concentration of purified human TREX1 protein together with the indicated concentration of purified human BAF protein. - = no TREX1 added. (B) Representative DNA gel from *in vitro* nuclease assay, using purified BAF K6A mutant with deficient DNA binding instead of wild type BAF in (A). (C) Representative DNA gel from *in vitro* nuclease assay, using purified human cGAS instead of wild type BAF in (A). (D) Quantification of the *in vitro* nuclease assay in (A)(B)(C); mean ± s.d., *n* = 3, Wilcoxon matched-pairs signed rank test between BAF and BAF K6A (**P<0.01). (E) Schematics of human TREX1 transgenes. FL = full length, TMD = trans-membrane domain, IDR = intrinsically disordered region. (F) Immunofluorescence of RPA in HeLa TREX1 KO cells stably expressing GFP-TREX1ΔTMD as shown in (E) and treated with CTRL or BAF siRNA. Arrows marked ruptured micronuclei indicated by the loss of NLS-RFP signal. Scale bar, 10 µm. (G) Quantification of normalized RPA intensity in ruptured micronuclei as shown in (F). Mean ± s.e.m., *n* = 3, Student’s *t*-test (****P < 0.0001), each experiment with greater than 100 total micronuclei quantified. See also Figure S5.

### ER-independent TREX1 resection in BAF-deficient cells

We previously reported that TREX1-ER tethering is critical for directing its activity against micronuclear DNA, but the mechanisms underlying ER dependency were poorly understood^8^. We reasoned that BAF accumulation at ruptured micronuclei may shield micronuclear DNA from the soluble TREX1ΔTMD protein and thus provide a mechanistic explanation for the critical nature of TREX1-ER tethering. To test this, we reconstituted *TREX1* KO cells with either wild-type GFP-TREX1 or a GFP-TREX1-ΔTMD mutant expected to lose its association with the ER (Figure 5E; Figure S5D). Consistent with prior reports^8,28^, GFP-TREX1-ΔTMD mutants exhibited a pancellular distribution that differed markedly from the ER co-localization of the wild-type GFP-TREX1 protein (Figure 5F). GFP-TREX1-ΔTMD mutants failed to accumulate at ruptured micronuclei and were completely defective for micronuclear DNA resection, as determined by measuring RPA32 signal intensity at ruptured micronuclei (Figures 5F and 5G). BAF depletion restored micronuclear DNA resection in GFP-TREX1-ΔTMD cells without promoting GFP-TREX1-ΔTMD localization to ruptured micronuclei (Figures 5F and 5G). Altogether, these data indicate that BAF shields micronuclear DNA from TREX1-mediated resection and suggest that ER tethering allows TREX1 to bypass BAF-mediated inhibition.

### BAF inhibits cGAS activation at ruptured micronuclei

BAF opposes cGAS activation by outcompeting cGAS for DNA binding within the nucleus^57^. However, it is unclear if BAF-dependent cGAS restriction extends to micronuclei, where cGAS exhibits a dramatic enrichment following micronuclear envelope rupture^5,6,8^. To test this, we performed immunofluorescence to measure cGAS signal intensity at ruptured micronuclei following BAF depletion. This analysis revealed a significant increase in cGAS levels at ruptured micronuclei in BAF-depleted cells suggesting that BAF competes with cGAS for binding to cytosol-exposed, micronuclear DNA (Figures 6A and 6B). To assess the impacts of BAF depletion on cGAS activation we used ELISA to measure cGAMP levels and qPCR to measure ISG expression following treatment with an inhibitor of the spindle assembly checkpoint kinase Mps1 to increase micronucleation rates^8^. In agreement with prior work^57^, BAF depletion led to a 3-fold increase in cGAMP levels (330 fmol cGAMP/mg protein in BAF-depleted cells vs 92 fmol cGAMP/mg protein in cells transfected with a control siRNA) potentially reflecting increased activation of the nuclear pool of cGAS (Figure 6C). *ISG54* and *ISG56* mRNA levels showed similar increases following BAF depletion (Figure 6D). In contrast to our prior work in chromosomally stable MCF10A cells^8^, *TREX1* deletion also significantly increased cGAMP levels in the absence of Mps1 inhibitor treatment, likely due to the high basal rates of micronucleation endogenous to HeLa cells (Figures 6C; Figures S6A). BAF-depletion led to a remarkable ∼10-fold increase in cGAMP levels in Mps1 inhibitor-treated cells relative to controls (1081 fmol cGAMP/mg protein in BAF-depleted, Mps1i-treated cells versus 133 fmol cGAMP/mg protein in Mps1i-treated controls). This effect was further increased 3-fold to 3107 fmol cGAMP/mg protein following BAF depletion in *TREX1* KO cells (Figures 6C). ISG expression followed similar patterns showing the strongest increases in Mps1i-treated cells deficient for BAF and TREX1 (Figure 6D). Thus, BAF and TREX1 work through complementary mechanisms to suppress immune responses in chromosomally unstable cells.

**Figure 6.**
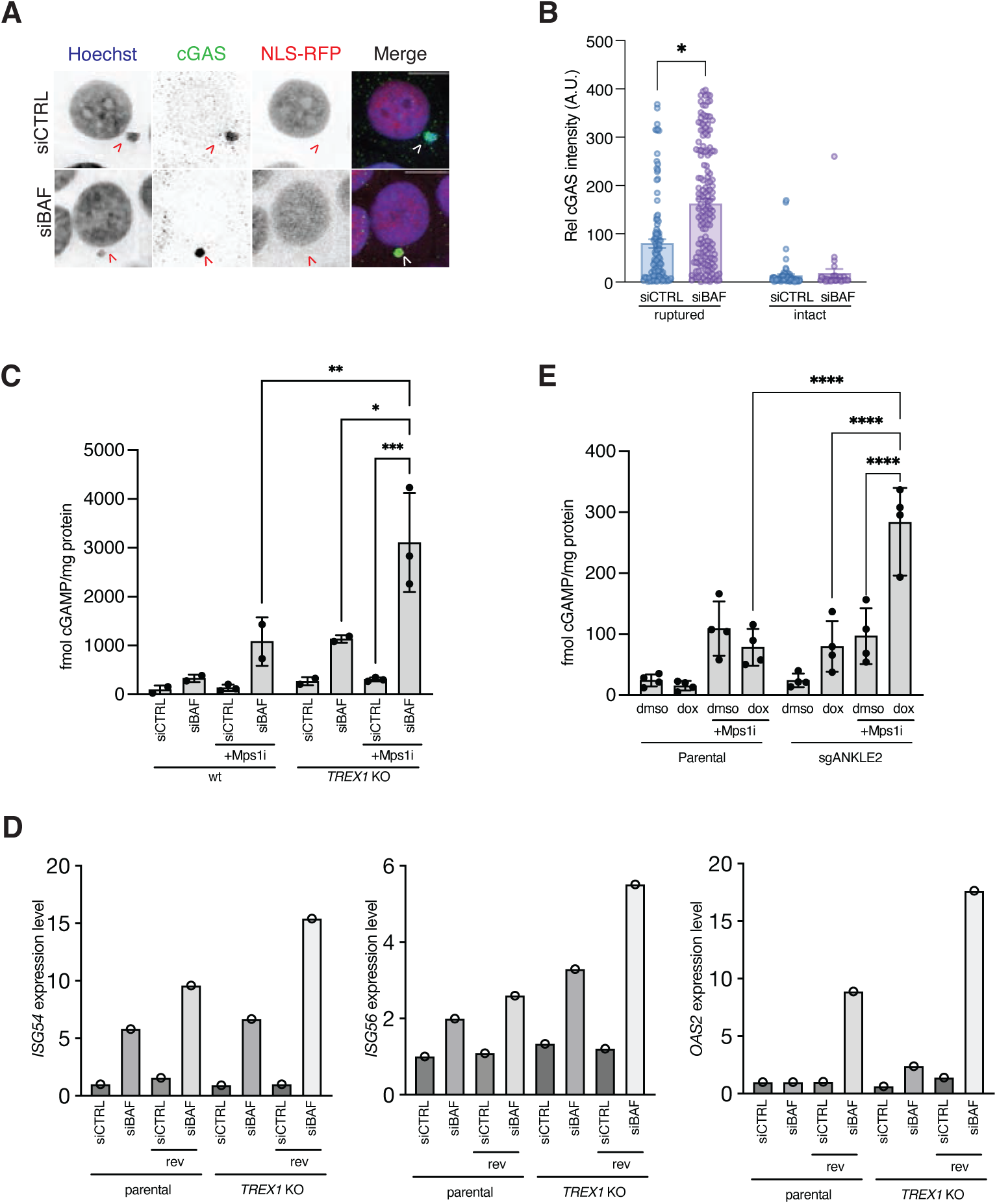
BAF inhibits cGAS activation at ruptured micronuclei. (A) Immunofluorescence of cGAS in HeLa iCas9 sgBAF expressing NLS-RFP treated with CTRL or BAF siRNA. Arrows marked ruptured micronuclei indicated by the loss of NLS-RFP signal. Scale bar, 10 µm. (B) Quantification of normalized cGAS intensity in ruptured micronuclei as in (A). mean ± s.e.m., *n* = 3, Student’s *t*-test (*P < 0.1), each experiment with greater than 100 total micronuclei quantified. (C) ELISA analysis of cGAMP production in the indicated cells. HeLa wt or TREX1 KO cells were treated with CTRL or BAF siRNA for 4 days. After 2 days of siRNA, cells were treated with 0.2µM Mps1 inhibitor reversine. (D) Quantitative PCR analysis of *ISG54*, *ISG56*, and *OAS2* transcripts in HeLa cells treated indicated siRNA for 4 days and reversine 0.2µM for 2 days. *n* = 1 experiment. (E) ELISA analysis of cGAMP production in the indicated cells. HeLa parental or sgANKLE2 cells were treated with dmso or dox for 4 days. Ond day post siRNA transfection, cells were treated with 0.5µM Mps1 inhibitor reversine for two days. For (C)(D), Mean and SD of *n*=3 experiments are shown. Ordinary one-way ANOVA with Dunnett’s multiple comparisons test (*P < 0.1, **P < 0.001, ***P < 0.001). See also Figure S6.

Phosphorylation of BAF by Vaccinia-related kinases diminishes its binding to DNA^58,59^. Recruitment of BAF to segregated chromosomes at mitotic exit depends on its dephosphorylation by the PP2A phosphatase, an activity promoted by the PP2A-interacting, LEM-domain protein Ankle2^60–62^. To determine if Ankle2-dependent dephosphorylation of BAF promotes its localization to ruptured micronuclei we performed immunofluorescence to monitor BAF subcellular localization in Ankle2-depleted cells. As expected, Ankle2 depletion resulted in slower migration of BAF in immunoblotting experiments, likely reflecting increased pools of phospho-BAF (Figure S6B). Reminiscent of prior observations at mitotic exit, Ankle2 depletion severely decreased BAF localization to ruptured micronuclei (Figures S6C and S6D). In contrast, depletion of the unrelated, LEM-domain protein LEMD2 had no apparent effect on BAF subcellular localization. In association with the BAF localization defect to ruptured micronuclei, Ankle2 depletion led to increased cGAS localization to ruptured micronuclei and a ∼4-fold increase in cGAMP levels following Mps1i treatment compared to controls (283 fmol cGAMP/mg protein in Ankle2-depleted, Mps1i-treated cells versus 73 fmol cGAMP/mg protein in Mps1i-treated controls) (Figures 6E; Figures S6E and S6F). Altogether, these data indicate that BAF DNA binding is critical for effective cGAS regulation at ruptured micronuclei.

## DISCUSSION

Here, we identify BAF as a critical barrier to DNA transactions comprising TREX1 resection and cGAS activation at ruptured micronuclei. In agreement with prior reports^4,43^, BAF accumulates at micronuclei following micronuclear membrane collapse. BAF localization to ruptured micronuclei depends on its DNA binding function indicating that BAF is likely recruited to ruptured micronuclei through direct attraction to cytosol-exposed genomic DNA. BAF concentrates additional factors at ruptured micronuclei, including LEM-domain proteins, such as Ankle2 and LEMD2, and TREX1-ER. Several lines of evidence suggest that BAF augments TREX1 recruitment to ruptured micronuclei via an indirect mechanism. First, TREX1 ER association is essential for its localization to ruptured micronuclei^8^. Second, TREX1 truncation mutations that remove most of the cytosolic N-terminal region— thus disrupting contact with cytosol-facing interactors—do not impair TREX1 localization to ruptured micronuclei^8^. Third, depletion of transmembrane LEM-domain proteins, including ER-localized Ankle2, phenocopy BAF in terms of diminishing TREX1 localization to ruptured micronuclei. Finally, co-immunoprecipitation experiments failed to detect an interaction between BAF and TREX1 (data not shown). Thus, BAF likely promotes timely TREX1 recruitment to ruptured micronuclei in an indirect mechanism whereby TREX1 localization depends on BAF-dependent concentration of ER membranes to ruptured micronuclei via its interaction with transmembrane LEM-domain proteins.

Despite promoting TREX1 recruitment to ruptured micronuclei, TREX1-dependent resection of micronuclear DNA is increased in BAF depleted cells. Moreover, TREX1 mediated resection of micronuclear DNA no longer required its C-terminal ER-tethering in the absence of BAF. BAF-mediated inhibition of TREX1 exonuclease activity is reminiscent of how BAF restricts cGAS activation by nuclear DNA^57^. However, while BAF inhibits cGAS through dynamic competition for DNA binding^57^, the mechanism of TREX1 inhibition appears to be more complex. BAF presents a barrier to soluble TREX1 and uses its DNA binding to antagonize TREX1 activity when larger concentration differences exist *in vitro*. TREX1 primarily engages the 3′ end of DNA substrates, but also makes extensive contacts with the non-substrate strand with 4 residues (F26, R128, K160, R164) contacting the DNA backbone^63^. BAF may destabilize TREX1-DNA interactions by competing with TREX1 for binding to the phosphate backbone of the non-substrate strand^64^. Of note, TREX1 K160 mutations are linked to autoimmune disease suggesting that TREX1 contacts with the non-substrate strand are likely critical for TREX1 function^64,65^. Alternatively, BAF may inhibit TREX1 DNA digestion through its DNA looping or compaction activities, which are thought to protect retroviral DNA by blocking retroviral integration machinery^50,66,67^.

While precise mechanisms require further investigation, BAF inhibits TREX1 resection at ruptured micronuclei and thus protects against micronuclear DNA damage. Mis-segregated chromosomes isolated within micronuclei undergo extensive DNA damage ultimately resulting in the catastrophic shattering phenomenon known as chromothripsis^54,68–72^. TREX1 activity promotes chromothriptic fragmentation in cellular models of telomere crisis^36,73^. BAF may therefore protect against the severity of chromothripsis rearrangements by suppressing TREX1 activity at ruptured micronuclei. Indeed, recent work indicates that interphase-associated micronuclear DNA damage may play a more modest role in chromothriptic fragmentation relative to a mitosis-specific chromosome shattering pathway driven by aberrant activation of the Fanconi Anemia pathway^74^.

Our results provide a mechanistic basis to explain TREX1’s reliance on ER tethering for micronuclear DNA resection and cGAS inhibition. As we previously reported^8^, TREX1 truncation mutants that are separated from the ER, such as TREX1ΔTMD, fail to localize to ruptured micronuclei and are unable to resect micronuclear DNA. Here, we show that BAF depletion unleashes TREX1ΔTMD activity at ruptured micronuclei thus suggesting that ER tethering is necessary because it allows TREX1 to bypass BAF-mediated inhibition. Although BAF’s opposing roles in promoting TREX1-ER recruitment and inhibiting TREX1 resection at ruptured micronuclei may appear paradoxical, these functions appear more coherent in light of BAF’s critical role during primary nuclear envelope rupturing^41,43^. While primary nuclear envelope repair is generally successful^42,75,76^, BAF protection of genomic DNA may protect against cGAS hyperreactivity and TREX1-mediated DNA damage in the interim period preceding membrane repair and restoration of compartmentalization. In the case of unsuccessful membrane repair events (e.g. ruptured micronuclei), BAF-mediated TREX1 recruitment may further shield against chronic cGAS activation, albeit at the expense of genome integrity. Linking to BAF, which has femtomolar affinity for DNA^50,55^, may help direct TREX1 to relevant targets throughout the cytosol and thus effectively compete with cGAS, which can exclude TREX1 from DNA through the formation of cGAS-DNA condensates^55^. Thus, BAF-mediated TREX1 recruitment may augment its ability to inhibit cGAS during cases of chronic DNA exposure, while BAF-mediated TREX1 shielding may preserve genome integrity during transient DNA exposure.

Original studies demonstrated delayed cGAS dependent responses to DNA damaging events^5,77^. Interferon stimulated gene expression became detectable at 48-72 hours after ionizing radiation and was maximal at six days despite the presence of micronuclei at earlier time points. Subsequent studies suggest requirements for micronuclear replication and entry into mitosis for cGAS dependent ISG expression^77,78^. Combined inhibition by BAF and TREX1 may thus explain delays in cGAS activation following DNA damage. Recent studies further argue that rapid cGAS activation at ruptured micronuclei in chromosomally unstable cells is held in check by histone-bound micronuclear DNA^79–83^. By showing that BAF suppresses cGAS activation at ruptured micronuclei, our data identify an additional regulatory layer against innate immune activation in chromosomally unstable cells. Unchecked TREX1 resection of micronuclear DNA partially compensates for BAF loss by imposing additional regulation of cGAS activation at ruptured micronuclei. Indeed, combined loss of BAF and TREX1 led to >20-fold increases in cGAMP levels in chromosomally unstable HeLa cells. These observations suggest that the presence of nucleosome-containing chromosomal DNA in micronuclei is insufficient to fully suppress cGAS activation. Interestingly, prior studies have noted that cGAS activation at micronuclei only occurs after a long delay following DNA damage in a manner that is dependent on mitotic progression^5,77^. In agreement, we observe greater than 10-fold increases in cGAMP production after MPS1i even in cells that express both BAF and TREX1, providing further evidence of cGAS activation within micronuclei. Future studies will be necessary to determine how mitotic progression and other events allow cGAS to bypass these regulatory mechanisms.

Taken together, our work identifies BAF as a critical gatekeeper of TREX1 resection and innate immune activation in chromosomally unstable cancer cells. Future work will be necessary to determine if BAF coating of micronuclear DNA may play important roles at other micronuclei-associated dysfunctions, such as transcriptional silencing^3,84^, histone post-translational modifications^85^, or DNA replication defects^69,86^.

## Limitations of the study

Our data indicate that BAF depletion delays, but does not completely eliminate TREX1 recruitment to ruptured micronuclei. Therefore, we expect that additional mechanisms, such as ESCRT-III-dependent membrane intercalation^39^, may be operative. Analysis of BAF L58R mutants together with Ankle2 and LEMD2 depletion phenotypes provide a general understanding of how BAF recruits TREX to membranes, but precise molecular details remain to be determined. Similarly, identifying the precise mechanism of BAF-mediated TREX1 inhibition will require further work.

## ACKNOWLEDGEMENTS

We thank members of the Maciejowski lab and L. Mohr for work on this manuscript. We thank K. Kulej from A.K’s laboratory for the help in uploading proteomic data. Work in J.M.’s laboratory is supported by the NCI (R37CA261183; R01CA270102; P30CA008748), the Pershing Square Sohn Cancer Research Alliance, the Frank A. Howard Scholars Program, the Mary Kay Ash Foundation and the Experimental Therapeutics Centers at MSKCC. Y.C. is supported by the EIO Scholars Program Fellowship from the Center for Experimental Immuno-oncology at MSKCC. A.K. is a scholar of the Leukemia & Lymphoma Society and is supported by NIH R01 CA204396 and P30CA008748. W.Z. receives support from the National Natural Science Foundation of China (NSFC) (32270920), the Shenzhen Basic Research Program (JCYJ20240813094802004), the Shenzhen Talent Program (KQTD20210811090115021), the Guangdong Innovative and Entrepreneurial Research Team Program (2021ZT09Y104), and the Shenzhen Science and Technology Program (ZDSYS20220402111000001).

## AUTHOR CONTRIBUTIONS

Conceptualization: Y.C. and J.M.; Methodology: Y.C. and J.M.; Investigation: Y.C., R.X.N., X.L., E.T., P.C., J.H., A.S.; Formal Analysis: Y.C., X.L., E.T.; Writing - Original Draft: Y.C., E.T., and J.M.; Writing - Review and Editing - all authors; Visualization: J.M., E.T., and Y.C..; Funding Acquisition: Y.C., A.K., W.Z., and J.M; Supervision: A.K., W.Z., and J.M.

## DECLARATION OF INTERESTS

J.M. reports patents pending for targeting the cGAS-STING pathway in cancer. A.K. is a consultant to Rgenta, Novartis, Blueprint Medicines, Syndax, and Sellas. The other authors declare no competing interests.

## STAR Methods

### KEY RESOURCES TABLE

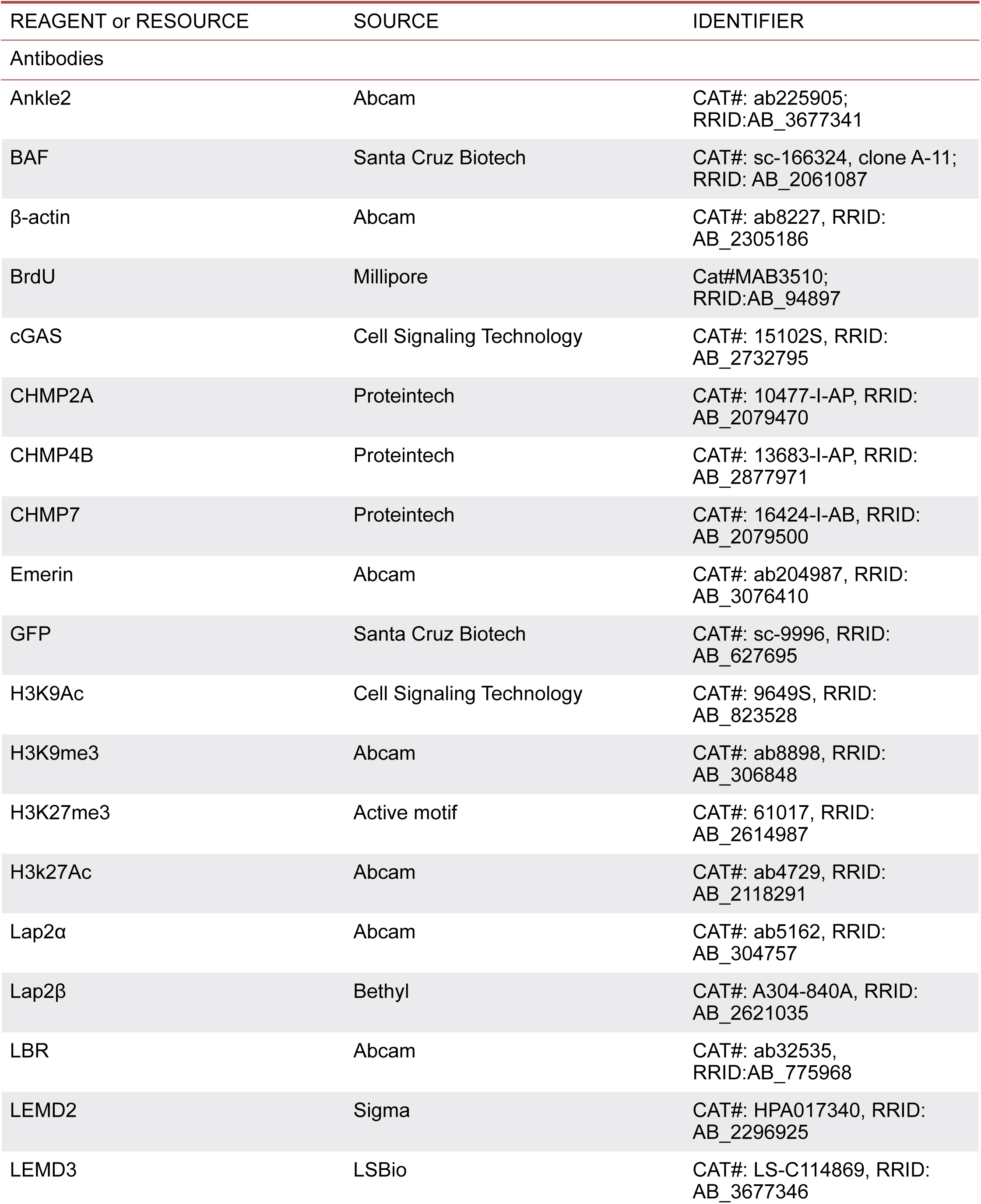
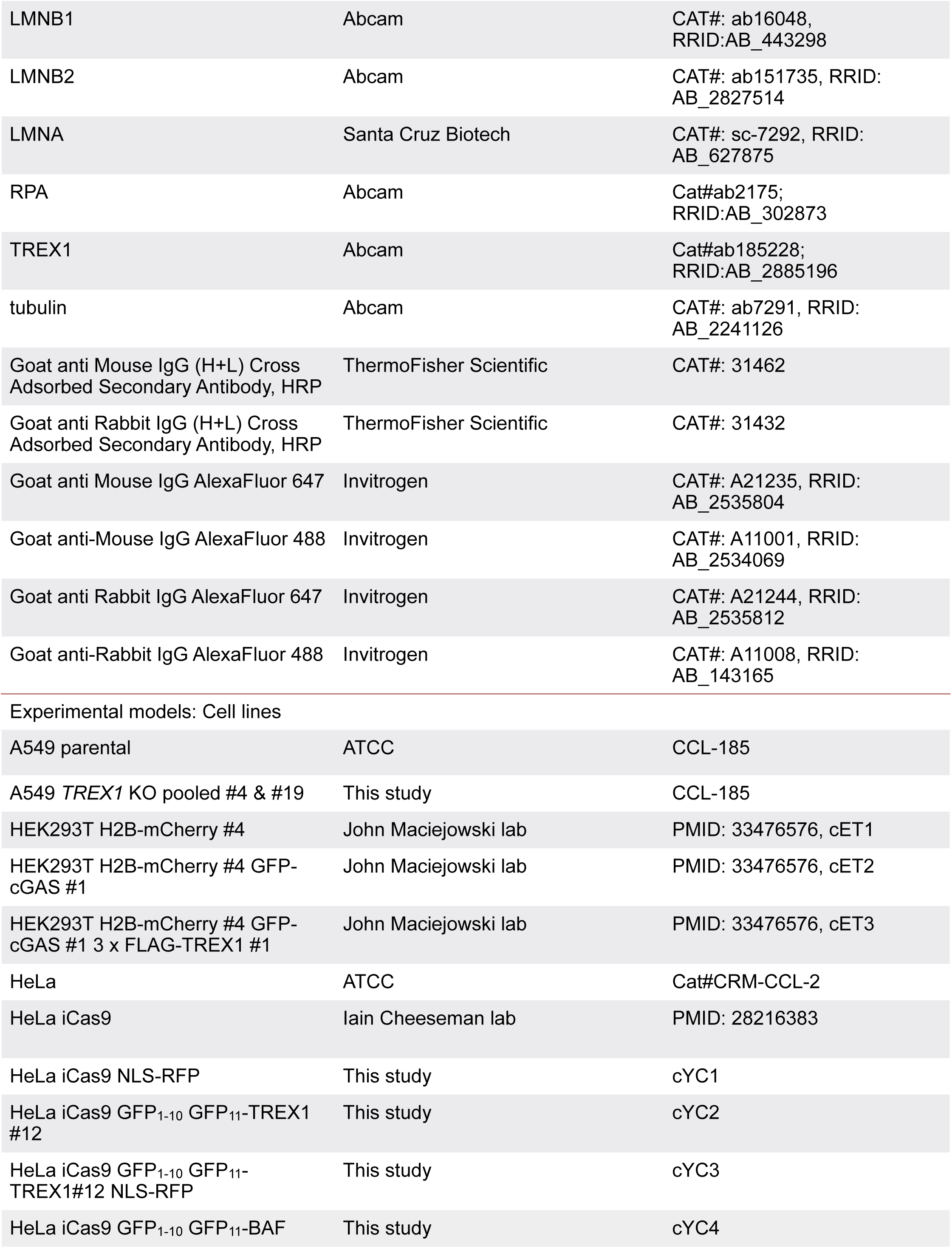
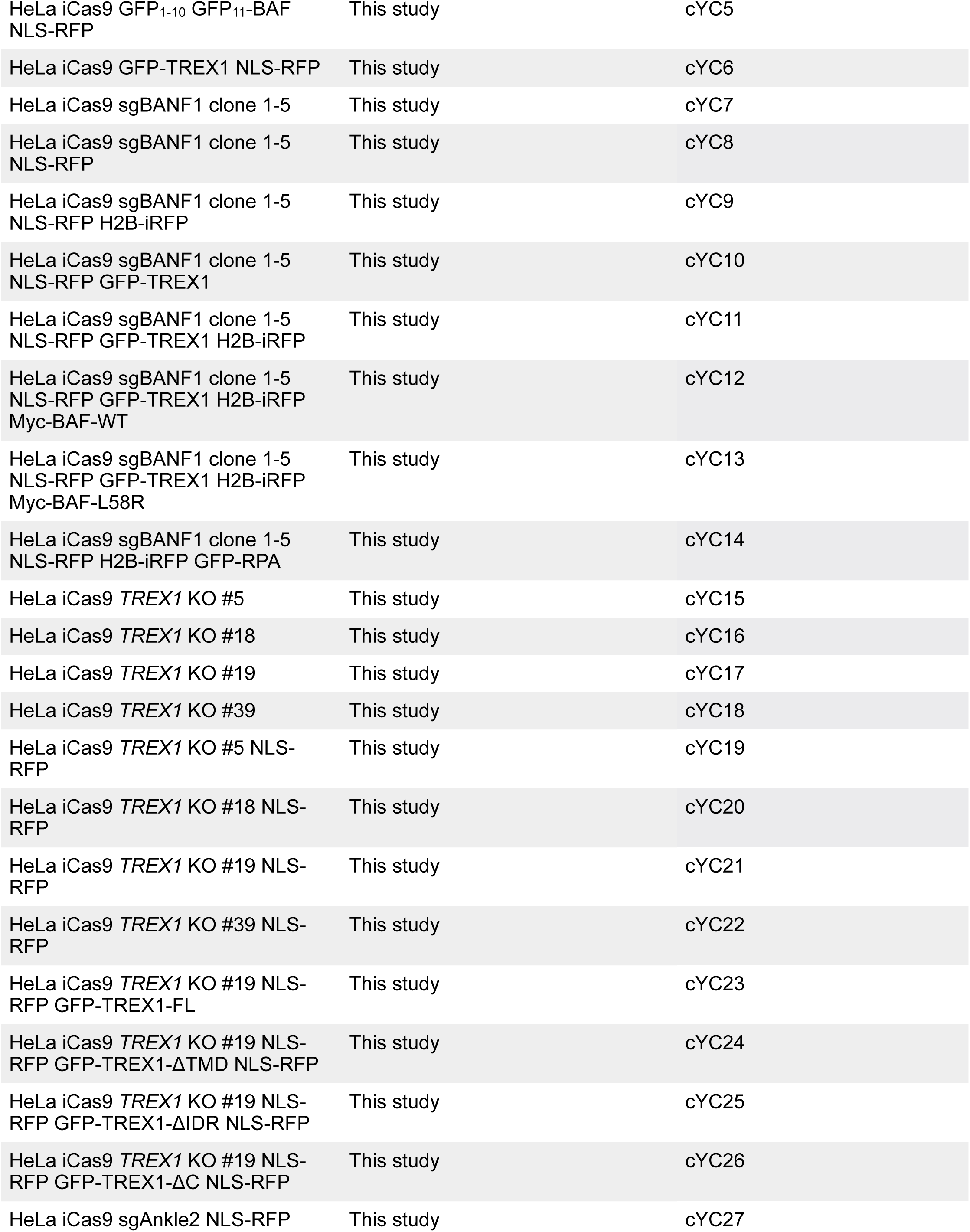
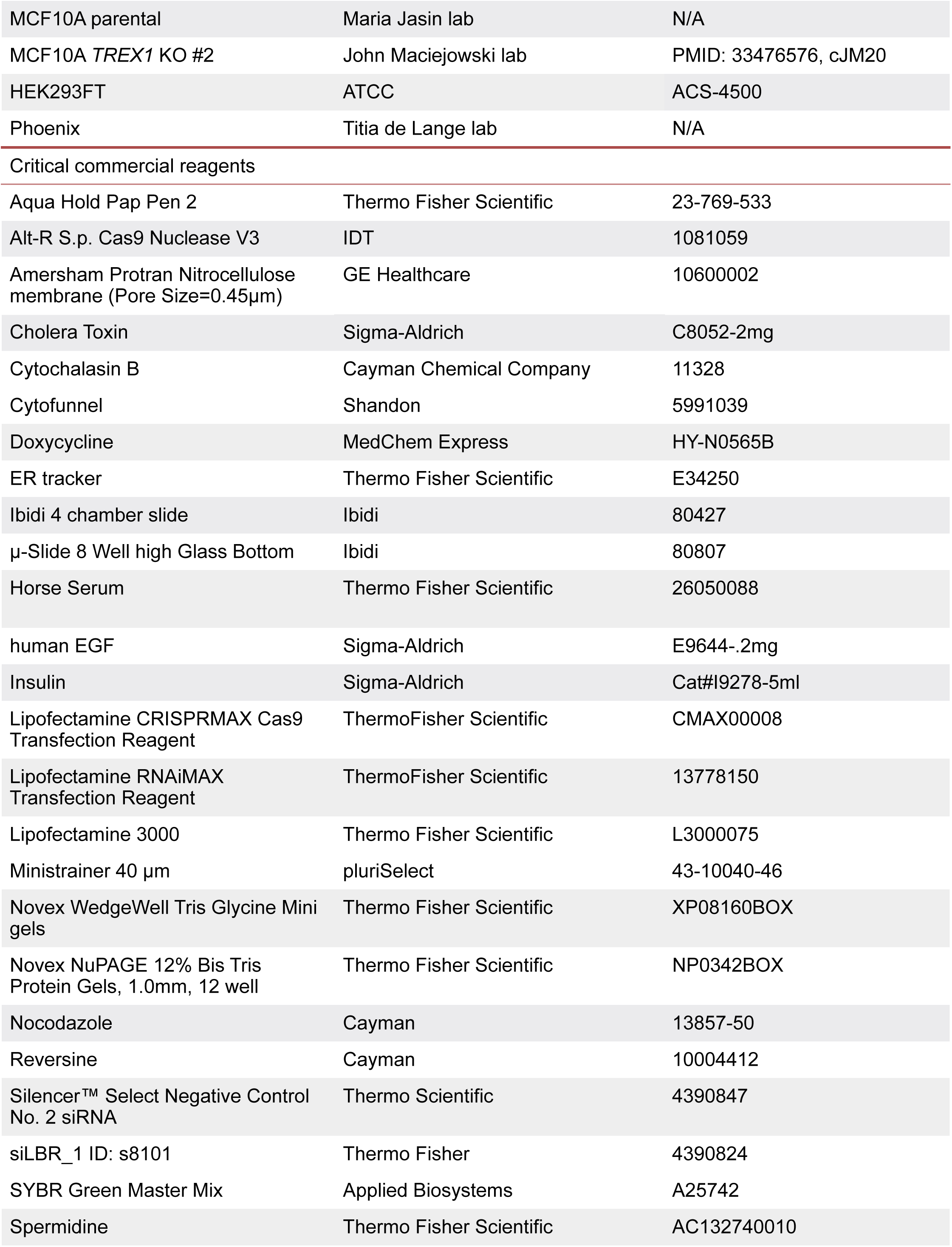
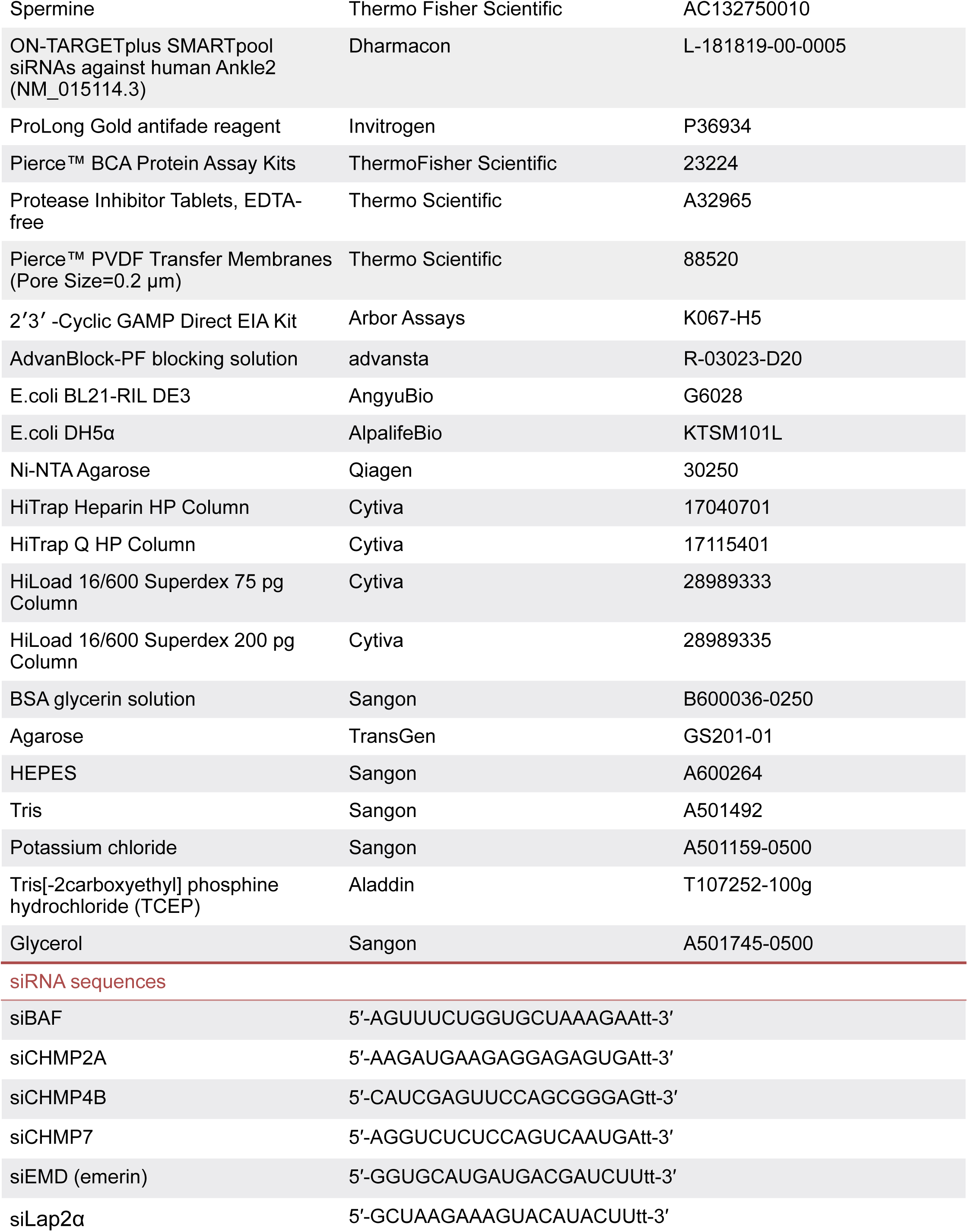
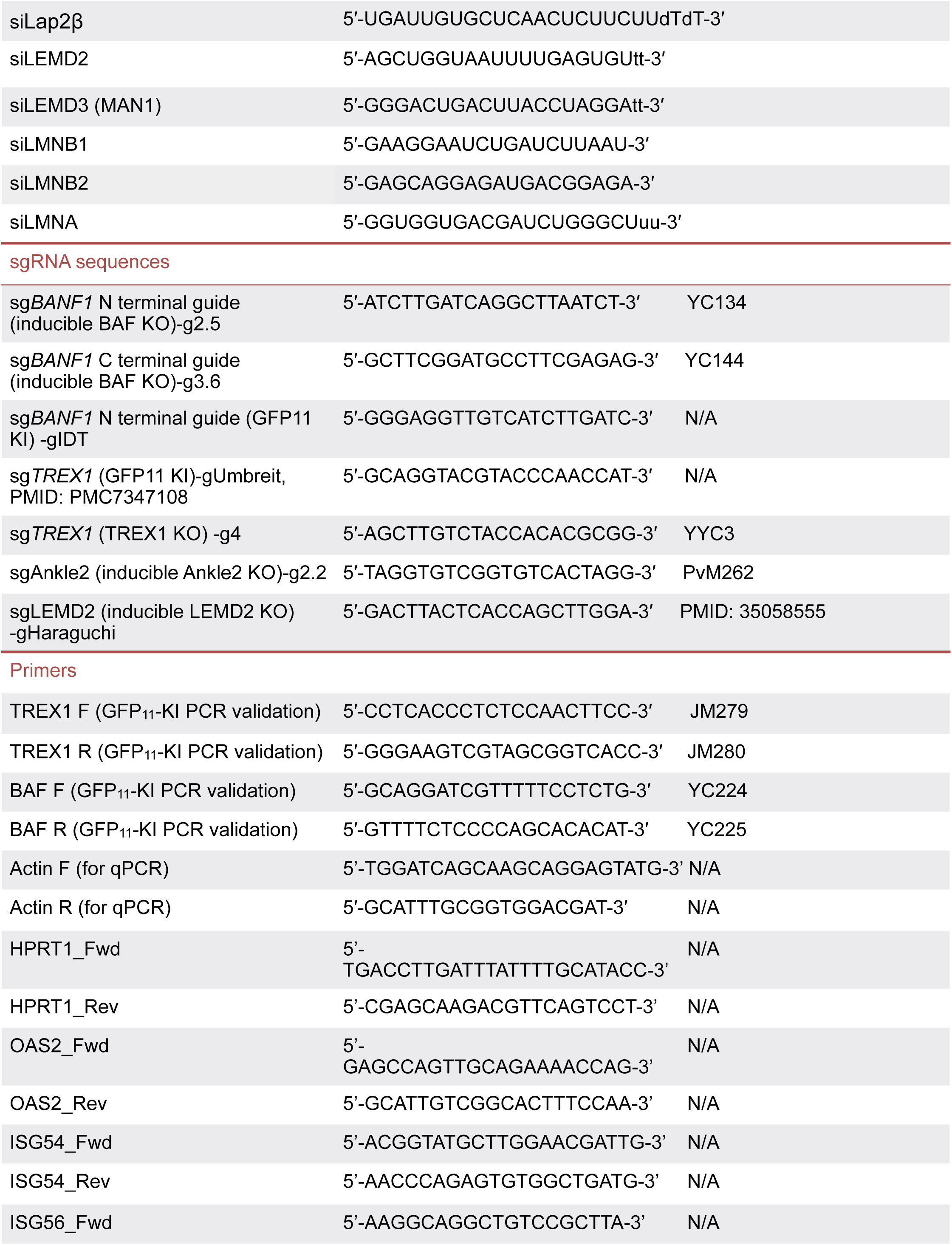
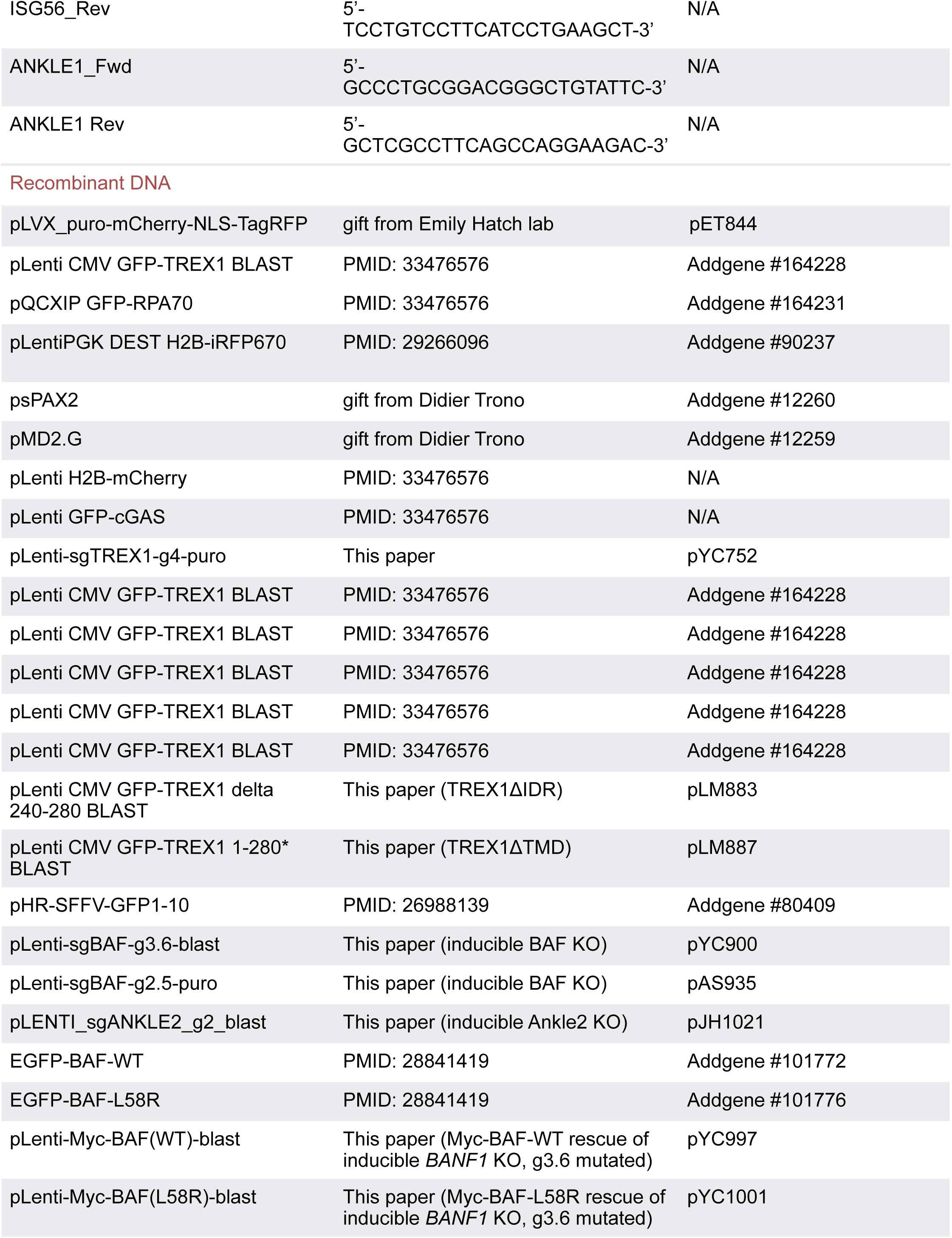
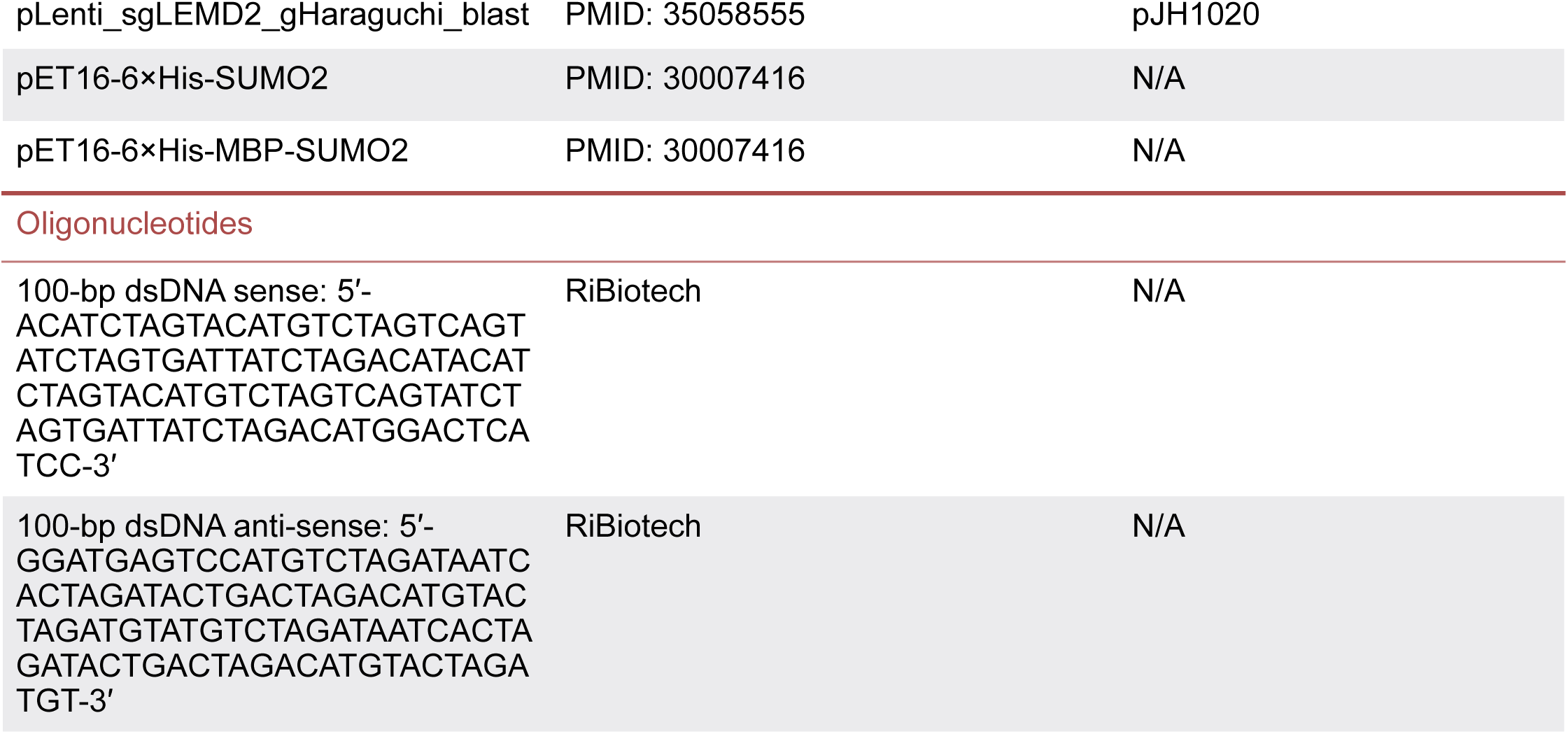

### RESOURCE AVAILABILITY

#### Lead contact

Further information and requests for resources and reagents should be directed to and will be fulfilled by the Lead Contact, John Maciejowski.

#### Materials availability

Cell lines generated in this study and listed in the Key Resource Table are available from Dr. John Maciejowski.

#### Data and code availability

The datasets and original source data generated during this study are available.

● All data reported in this paper will be shared by the lead contact upon request.
● The paper does not report original code.

### EXPERIMENTAL MODEL DETAILS

#### Cell culture

HeLa, HeLa iCas9, HEK293T, Phoenix, 293FT were grown in DMEM supplemented with 10% FBS. MCF10A cells were cultured in 1:1 mixture of F12:DMEM media supplemented with 5 % horse serum (Thermo Fisher Scientific), 20 ng/mL human EGF (Sigma), 0.5 mg/mL hydrocortisone (Sigma), 100 ng/mL cholera toxin (Sigma) and 10 µg/mL recombinant human insulin (Sigma). A549 (ATCC) cells were cultured in DMEM supplemented with 10 % FBS and 1% glutamine. All media was supplemented with 1% penicillin-streptomycin. Unless otherwise noted, all media and supplements were supplied by the MSKCC core facility. HeLa iCas9 is a gift from Iain Cheeseman lab from MIT.

#### Generation of TREX1 KO cell lines

For HeLa, *TREX1* KO cells were generated by stably expression of TREX1 gRNA (Key Resource Table) in HeLa iCas9 cells followed by 1 µg/mL puromycin selection. HeLa iCas9 *TREX1* KO polyclonal population was treated with 1 µg/mL doxycycline for 3 days, followed by limited dilution. HeLa iCas9 *TREX1* KO clones were selected by immunoblotting for TREX1 with extended exposure to ensure thorough depletion of TREX1. For A549, *TREX1* KO cells were generated by lipofection (Lipofectamine3000) of TREX1 gRNAs cloned in Cas9-gRNA-all (with mCherry) in one plasmids (Addgene #64324) followed by FACS selection of mCherry-positive cells. *TREX1* KO polyclonal population was subcloned by limiting dilution and *TREX1* KO clones were selected by immunoblotting. MCF10A *TREX1* KO was generated as previously described^8^.

#### Generation of inducible *BANF1* KO

A few gRNAs were designed using Benchling to target the start and the end of the *BANF1* coding region (Supplementary Figure S2J). Guide RNAs targeting the beginning of *BANF1* coding region were cloned into pLenti-Guide-Puro (Addgene #52963), and guide RNAs targeting the end of *BANF1* coding region were cloned into pLenti-Guide-Blast (Addgene #199622). HeLa iCas9 parental cells (Cas9 under a tetracycline-inducible promoter, a gift from Iain Cheeseman lab from MIT) were stably expressed with a combination of start and end gRNAs and selected with 1 µg/mL puromycin and 5 µg/mL blasticidin. The polyclonal populations (from different gRNA combinations) were treated with 1µg/mL doxycycline for 72 hours (to induce Cas9 expression) and compared to the BAF protein level of the cells treated with dmso. The gRNA combination with the most BAF protein depletion was chosen, and the polyclonal population was further subcloned by limiting dilution and a single clone was chosen based on its depletion level (comparable to BAF siRNA depletion as in (Figure S2K).

#### Endogenously tagged TREX1 and BAF by split GFP system

Using a split GFP system^87^, GFP_1-10_ (Addgene plasmid #80409) was stably expressed in HeLa iCas9 parental cells and CRISPR/Cas9-mediated HDR was performed to knock in GFP_11_-3×FLAG at the N-terminus of TREX1 (Figure 2A). Cas9: sgRNA of 1:1 molar ratio was incubated for 15-20 minutes at room temperature to assemble RNP. 1 million cells were nucleofected together with ssDNA ODN template (100 pmol) by Lonza 4D nucleofector. ssDNA ODN template encompasses 70 bp homology arms and GFP_11_-3×FLAG and was synthesized by IDT technologies. CRISPR knockin was performed in HeLa iCas9 GFP_1-10_ cells and GFP-positive clones were selected by FACS. GFP positive clones were subcloned by limiting dilution and confirmed by PCR and western blot. While most TREX1 alleles are tagged with GFP_11_-3×FLAG, a small proportion of unmodified TREX1 persisted due to the polyallelic nature of HeLa cell lines (Figure 2B).

For BAF, GFP_11_-3×FLAG-miniAID was knocked into the N-terminus of BAF in HeLa iCas9 cells that stably express GFP_1-10_. Cas9: sgRNA of 1:1 molar ratio was incubated for 15-20 minutes at room temperature to assemble RNP. 1 million cells were nucleofected together with 2.5 µg donor plasmid by Lonza 4D nucleofector. Donor plasmid was composed of pUC19 base vector (Addgene #50005), GFP_11_-3×FLAG-mAID, and 500 bp homology arms (Figure S1E). Successful knock-in cells were enriched through two rounds of FACS sorting for GFP-positive cells and used as polyclonal populations.

#### Reconstituted GFP-TREX1 mutants

TREX1 ER-tethering mutant has a truncated transmembrane domain (TREX1ΔTMD) within the amino acid 285 to 314 (Figure 5E). Unlike TREX1ΔC, which removes the entire C-terminal including the intrinsically disordered region (IDR), TREX1ΔTMD retains the IDR. Additionally, TREX1-FL was generated as controls. Open reading frames were cloned into pLenti-CMV-GFP plasmid (Addgene #164228). Constructs were transfected into HEK293FT cells together with psPAX2 (Addgene #12260) and pMD2.G (Addgene #12259) using calcium phosphate precipitation. Supernatants containing lentivirus were filtered and supplemented with 4 µg/mL polybrene. Successfully transduced cells were selected by FACS of GFP-positive cells. The absence of endogenous TREX1 and the expression of the transgenes were validated by immunoblotting (Figure S5D). TREX1-FL mutants showing autoclipping and forming bands of the same size as endogenous TREX1.

#### Pharmacological treatments to generate micronuclei

To induce micronucleation in siRNA screen (Figure 2E), immunofluorescence (Figures 4C, E, J, and I), and live cell imaging (Figures 2H, 3A, 3F, and 4A) experiments, cells were treated with 100 ng/mL nocodazole for 6 hours followed by five times of media wash the day before live cell imaging or fixation as previously described^4,86^.

To induce micronucleation in cGAMP-ELISA experiments (Figure 6C), cells were treated with 4 days of siRNA to deplete BAF, and treated with 0.2 µM reversine for 2 days before harvest (within 4 days of siRNA treatment). To induce micronucleation in cGAMP-ELISA of Ankle2 depleted cells (Figure 6E), cells were treated total 4 days of dmso or dox (1µg/mL), and treated with 0.5 µM reversine for 3 days before harvest (within 4 days of dmso/dox treatment).

#### Immunofluorescence

For immunofluorescence microscopy of micronucleated cells, HeLa cells, seeded on coverslips 24 hours before, were treated with siRNA (Figure 4E, Figure S4B, D, F, and H) or doxycycline 1 µg/mL for 72 hours (Figure 4C, G, I; Figure 6B) . The day before fixation, cells were treated with 100 ng/mL nocodazole for 6 hours followed by five times of media wash. After the treatment, cells were carefully washed with PBS prior to fixation in 2 % paraformaldehyde in PBS for 12 minutes. Coverslips were washed with TBS, incubated in TBS with 0.5 % Triton X-100 for 5 minutes and washed again with TBS. Coverslips were incubated in blocking buffer (1 mg/ mL BSA, 3 % goat serum, 0. 1% Triton X-100, 1 mM EDTA in PBS) for 1 hour and incubated in primary antibodies (see Key Resources Table) diluted in blocking buffer for 2 hours at room temperature. After 3 washes with TBS-TX (0.1 % Triton X-100), coverslips were incubated with secondary antibodies (see Key Resources Table), diluted in blocking buffer, for 1 hour, then washed 3 times with TBS-TX. DNA was stained with DAPI or Hoechst (both at 1 µg/mL) for 10 minutes, before coverslips were washed with TBS. Coverslips were mounted in ProLong Gold Antifade Mountant (Life Technologies).

Immunofluorescence images were acquired on a Nikon Eclipse Ti2-E equipped with a CSU-W1 spinning disk with Borealis microadapter, Perfect Focus 4, motorized turret and encoded stage, polycarbonate thermal box, 5 line laser launch [405 (100 mw), 445 (45 mw), 488 (100 mw), 561 (80 mw), 640 (75 mw)], PRIME 95B Monochrome Digital Camera and 100× 1.45 NA objective. Images were further analyzed in Fiji.

Working concentrations for immunofluorescence: Anti-Ankle2 (1:100, Abcam, ab225905), Anti-BrdU (1:500, Millipore, MAB3510), Anti-cGAS (1:500, Cell Signaling Technology, 15102S), Anti-CHMP4B (1:100, Proteintech, 13683-I-AP), Anti-CHMP7 (1:100, Proteintech, 16424-I-AB), Anti-emerin (1:200, Abcam, ab204987), Anti-H3K9Ac (1:400, Cell Signaling Technology, 9649S), Anti-H3K9me3 (1:500, Abcam, ab8898) Anti-H3K27me3 (1:1000, Active Motif, 61017), Anti-H3K27Ac (1:200, Abcam, ab4729), Anti-LEMD2 (1:100, Sigma, HPA017340), Anti-LMNB1 (1:100, Abcam, ab16048), Anti-LMNA (1:250, Santa Cruz Biotech, sc-7292) Anti-RPA (1:250, Abcam, ab2175).

#### Native BrdU staining

For detection of ssDNA in MN, cells seeded on coverslips were preincubated with BrdU (10 mM) for 48 hours before fixation for 24 hours. BrdU was washed out with media twice and treated with nocodazole (100 ng/mL) for 6 hours and then washed with media for 5 times before fixation. Before BrdU immunofluorescence, cells were carefully washed with PBS, incubated in ice cold extraction buffer (10 mM Pipes-NaOH pH 7, 100 mM NaCl, 300 mM sucrose, 3 mM MgCl_2_, 1 mM EGTA, 0.5 % Triton X-100) for 10 minutes. Fixation and staining steps, as well as quantification were performed as described above.

#### Image analysis

Image analysis was performed in ImageJ.

For the siRNA screen (Figure 2E), ruptured MN with loss of NLS-RFP signal were manually scored as GFP-TREX1 positive or negative GFP-TREX1 ruptured MN. Percentage of GFP-TREX1 positive ruptured MN were calculated with more than 100 total micronuclei analysed each repeat, at least three independent experiments repeated for each siRNA target.

Quantification of GFP-TREX1 MN/ER ratio (Figure 2H, 3F), MN were identified by H2B-iRFP signal. ROI was manually drawn in the center of MN and neighboring ER, and GFP-TREX1 intensities at MN and ER are measured. GFP-TREX1 MN/ER ratios were calculated after removal of background signal.

Quantification of the ER tracker signal (Figure 3A), MN were identified by H2B-iRFP signal. ROI was manually drawn in the center of MN and ER tracker signals were measured and then removed background signals. Quantification of GFP-RPA foci in Figures (Figure 4A), MN were identified by H2B-iRFP signal. MN were manually called RPA32-positive with equal or more than 4+ RPA32 foci present in MN. Fractions of RPA+ MN were calculated at each time point.

Quantification of RPA32-, γH2AX- or BrdU-MN intensity was performed as follows (Figure 4C,E,J,I): MN were identified by Hoechst or DAPI signal, then the NLS-RFP signal was used to distinguish between intact (NLS-RFP-positive) and ruptured MN (NLS-RFP-negative). ROI was manually drawn in the center of MN and intensities of proteins of interest were measured.

#### Immunoblotting

For immunoblotting in Supplementary Figure S2B, S2E, S3F, 1 × 10^6^ cells were harvested by trypsinization and lysed in RIPA buffer (150 mM NaCl, 50 mM Tris-HCl, pH 8, 1 % NP-40, 0.5 % sodium deoxycholate, 0.1 % SDS) supplemented with 0.5 mM PMSF and Pierce Protease Inhibitor Tablet, EDTA free. Cells were shaken at 4 °C 800 × *g* in RIPA for 30 minutes, and lysates were sheared through 28 ½-gauge syringes 10 times until the sample is smooth. Lysates were cleared by centrifugation at 20,000 × *g* for 20 minutes at 4 °C, and the protein concentration was quantified using the Pierce BCA Protein Assay kit (ThermoFisher Scientific). 30 µg of protein was loaded into Tris-Glycine gels (Thermo Fisher Scientific). Protein transfer was performed by wet transfer with 1X Towbin Buffer (25 mM Tris, 192 mM glycine, 0.01 % SDS, 20 % methanol) and nitrocellulose membranes at 100 V for 1 hour on ice. Membranes were blocked in 1X AdvanBlock-AF (Protein Free) blocking solution (advansta) for 1 hour at room temperature and incubated with primary antibody (See Key Resources Table) diluted in AdvanBlock-Protein Free blocking solution (concentration see below) overnight at 4 °C. Membranes were washed four times in 1X TBS-T followed by incubation with horseradish-peroxidase (HRP)-conjugated secondary antibodies (See Key Resources Table) diluted 1:10,000 in blocking buffer for 1 hour at room temperature. After four washes in 1X TBS-T, membranes were rinsed in TBS and visualized using enhanced chemiluminescence (ThermoFisher Scientific).

For immunoblotting of BAF shown in Supplementary Figure S2I, S2K, 50 µg protein was loaded on 12 % Bis-Tris protein gels (Invitrogen, NP0342BOX). Precision Plus Protein™ Dual Color Standards (Bio-Rad) was used to visualize BAF at 10 kDa. Gels were transferred to 0.2 µm PVDF membrane (Thermo Fisher) at 50 V for 1 hour on ice. Membranes were blocked in 1X AdvanBlock-AF blocking solution (advansta) for 1 hour at room temperature and incubated with primary antibody (Santa Cruz, sc-166324) diluted 1:500 in blocking solution overnight at 4 °C. Membranes were washed four times in 1X TBS-T followed by incubation with horseradish-peroxidase (HRP)-conjugated secondary antibodies diluted 1:10,000 in blocking buffer for 1 hour at room temperature. After four washes in TBS-T, membranes were detected with Pierce ECL Western Blotting Substrate (Thermo Scientific) supplemented with 10 % Lumigen ECL Ultra (Lumigen) and imaging was performed using enhanced chemiluminescence (Amersham Image Quant 800 and BioRad ChemiDoc MP). For immunobloting of purified micronuclei (Figure 1H), samples were lysed in 1× Laemmli buffer at 2.5 × 10^7^ micronuclei/mL. Lysates were denatured at 95 °C for 5 minutes and DNA was sonicated using a Bioruptor 300 (Diagenode) on the high setting for 8 cycles 30 sec ON / 30 sec OFF. Lysate equivalents to 0.5 × 10^6^ micronuclei were loaded in each lane and processed as described above.

Working concentrations for immunoblotting:

Anti-Ankle2 (WB, 1:500, Abcam, ab225905), Anti-BANF1 (WB, 1:500, Santa Cruz Biotech, clone A-11, sc-166324), Anti-β-actin (WB, 1:2000, Abcam, ab8224), Anti-cGAS (WB, 1:500, Cell Signaling Technology, 15102S), Anti-CHMP2A (WB, 1:500, Proteintech, 10477-I-AP), Anti-CHMP4B (WB, 1:1000, Proteintech, 13683-I-AP), Anti-CHMP7 (WB, 1:1000, Proteintech, 16424-I-AB), Anti-emerin (WB, 1:500, Abcam, ab204987) Anti-GFP (WB, 1:1000, Santa Cruz Biotech, sc-9996), Anti-Lap2α (WB, 1:1000, Abcam, ab5162), Anti-Lap2β (WB, 1:1000, Bethyl, A304-840A), Anti-LBR (WB, 1:1000, Abcam, ab32525), Anti-LEMD2 (WB, 1:500, Sigma, HPA017340), Anti-LEMD3 (WB, 1:1000, LSBio, LS-C114869), Anti-LMNB1 (WB,1:500, Abcam, ab16048) Anti-LMNB2 (WB, 1:500, Abcam, ab151735), Anti-LMNA (WB, 1:500, Santa Cruz Biotech, sc-7292), Anti-TREX1 (WB, 1:1000, Abcam, ab185228), Anti-tubulin (WB, 1:2000, Abcam, ab7291).

#### Lentivirus and retrovirus production and infection

For lentiviral transduction, constructs were transfected into 293FT cells together with psPAX2 (Addgene #12260) and pMD2.G (Addgene #12259) using calcium phosphate precipitation. Supernatants containing lentivirus were filtered and supplemented with 4 µg/mL polybrene. Successfully transduced cells were selected using Puromycin (Fisher, 1 µg/mL) or Blasticidin (Fisher, 5 µg/mL), or flow sorting. Clones were isolated by limiting dilution.

For retroviral transduction, constructs were transfected into Phoenix amphotropic packaging cells using calcium phosphate precipitation. Cell supernatants containing retrovirus were filtered, mixed 1:1 with target cell media and supplemented with 4 µg/mL polybrene. Successfully transduced cells (GFP-RPA) were selected using flow sorting.

#### Live-cell imaging

For time-lapsed live-cell imaging, cells were treated with siRNA or doxycycline for 72 hours and treated with 100 ng/mL nocodazole for 6 hours the day before imaging, as described above for BAF depletion and MN enrichment. Live-cell imaging was performed at 37 °C using Nikon Eclipse Ti2-E equipped with CSU-W1 SoRa spinning disk super resolution confocal system, Borealis microadapter, Perfect Focus 4, motorized turret and encoded stage, 5-line laser launch [405 (100 mw), 445 (45 mw), 488 (100 mw), 561 (80 mw), 640 (75 mw)], PRIME 95B Monochrome Digital Camera, and CFI Apo TIRF 60x 1.49 NA objective lens. Cells were imaged in 37 °C and 5 % CO_2_. Images were acquired using NIS-Elements Advanced Research Software on a Dual Xeon Imaging workstation, with 10 minutes interval for 12 hours. Images containing ruptured MN were cropped and analyzed in Fiji. For testing ER localization upon BAF depletion, 1 µM ER tracker (Invitrogen) was added an hour before imaging and cells were stained for 30 minutes and then washed and replenished with FluoroBrite media.

#### siRNA screen

For siRNA screen, cells were transfected the day after seeding with the indicated siRNA together with Lipofectamine RNAiMAX (Invitrogen) (Key Resources Table) as instructed by the manufacturer’s protocol. All siRNA were used with final 100 nM concentration for a total of 72 hours, except CHMP2A (50 nM for 48 hours), CHMP4B and CHMP7 (50 nM for 48 hours serial transfection), as previously described in Vietri et al. 2020. siControl (Thermo Fisher) was transfected to serve as comparison. Cells were treated with 100 ng/mL nocodazole for 6 hours and washed out 5 times with media the day before live-cell imaging. At 72 hours of treatment, cells were stained with Hoechst for 1 ug/mL in Fluorobrite (Gibco) media supplemented with 10 % FBS and 1 % Pen/Strep. Live-cell imaging was performed on Hoechst-stained cells with a Nikon Eclipse Ti2-E as described in the immunofluorescence section. Images were analyzed with Fiji and percentage of GFP11-TREX1+ MN were analysed by manually examining each micronucleus to call for GFP positive or negative.

#### 2′3′-cGAMP quantification

0.3 × 10^6^ of HeLa cells were seeded into 6-cm dishes, and 24 hours later cells were treated with CTRL or BAF siRNA (100 nM) with lipofectamine RNAiMAX. 2 days after siRNA treatment, cells were treated with dmso or reversine (0.2 µM) for another 2 days (total 4 days of siRNA treatment), then harvested for cGAMP-ELISA treatment. For stimulation with HT-DNA, 1 × 10^6^ of HeLa cells were seeded into 10-cm dishes, and treated with dmso or dox for protein depletion (sgAnkle2) for a total of 72 hours. 24 hours before harvest, cells were transfected with 4 µg HT-DNA using Lipofectamine 3000 transfection reagent (Invitrogen) per manufacturer’s instructions. 24 hours post transfection, cells were washed with PBS, pelleted, and stored at -80 °C. To quantify 2′3′-cGAMP levels, 4 × 10^6^ cells were thoroughly resuspended in 120 µL lysis buffer (20 mM Tris-HCl pH 7.7, 100 mM NaCl, 10 mM NaF, 20 mM

β-glycerophosphate, 5 mM MgCl_2_, 0.1 % Triton X-100, 5 % glycerol) and lysed with a 28 ½ gauge needle. Lysates were incubated on ice for 30 minutes, centrifuged at 16,000 × *g*, 4 °C for 10 minutes and 2′3′-cGAMP levels were quantified using the 2′3′-cGAMP ELISA Kit (Arbor Assays) according to the manufacturer’s instructions.

#### *in vitro* nuclease assay of recombinant TREX1 and BAF

*In vitro* DNA degradation assays were performed as described previously with minor modifications (Zhou et al. 2022). Briefly, 1 µM of a 100-bp dsDNA substrate (sequence provided below) was incubated with 0.1 µM human or fish (*Labeo rohita*) TREX1, along with 0.1–10 µM of wild-type or DNA-binding-deficient (K6A) human BAF, in a 20 µL reaction mixture (20 mM Tris–HCl pH 7.5, 15 mM NaCl, 135 mM KCl, 5 mM MgCl_2_, and 1 mg mL-1 BSA) at 25 °C for 30 minutes. Reactions were quenched with SDS (final concentration 0.0167 % w/v) and EDTA (final concentration 10 mM), followed by incubation at 75 °C for 15 minutes. The remaining DNA was separated on a 4 % agarose gel in 0.5 × TB buffer (45 mM Tris, 45 mM boric acid). After electrophoresis, gels were stained in 0.5x TB containing 10 µg/mL ethidium bromide at 25 °C for 15 minutes, then destained in milli-Q water for 45 minutes. DNA was visualized by ImageQuant 800 Imaging System and quantified using FIJI software (Schindelin et al., Nat. Methods 2012; PMID 22743772).

100-bp dsDNA sense:

5′-ACATCTAGTACATGTCTAGTCAGTATCTAGTGATTATCTAGACATACATCTAGTACATGTCTAGTCAGTATCTAGTGATTATCTAGACATGGACTCATCC -3′

100-bp dsDNA anti-sense:

5′-GGATGAGTCCATGTCTAGATAATCACTAGATACTGACTAGACATGTACTAGATGTATGTCTAGATAATCACTAGATACTGACTAGACATGTACTAGATGT -3′

#### Protein expression and purification

Large-scale protein expression was performed as previously described^55,88^. Briefly, the DNA sequences of recombinant proteins were cloned into a custom pET16 vector for expression of a 6×His-SUMO2 fusion protein in E. coli BL21-RIL DE3 bacteria (Agilent) co-transformed with a pRARE2 tRNA plasmid. Starter cultures of E. coli were grown in MDG media, subsequently cultured in ∼2 L of M9ZB media, and induced with IPTG. Bacterial cultures were pelleted, flash-frozen in liquid nitrogen, and stored at −80 °C until purification. Protein purification was performed as previously described^55,88,89^. Briefly, bacterial pellets were resuspended in lysis buffer (20 mM HEPES-KOH pH 7.5, 400 mM NaCl, 10 % glycerol, 30 mM imidazole, 1 mM DTT) and lysed by sonication. The initial purification was performed using Ni-NTA (QIAGEN) affinity chromatography. Protein eluted from Ni-NTA was supplemented with ∼250 µg of human SENP2 protease to remove the SUMO2 solubility tag and dialyzed in dialysis buffer (20 mM HEPES-KOH pH 7.5, 150 mM NaCl, 1 mM DTT) at 4 °C for ∼14 hours. Untagged protein was further purified using Heparin HP ion-exchange (GE Healthcare) and eluted with a gradient of 150–1000 mM NaCl. Target protein was then further purified with size-exclusion chromatography using a 16/600 Superdex S75 column (GE Healthcare) equilibrated with protein storage buffer (20 mM HEPES-KOH pH 7.5, 250 mM KCl, 1 mM TCEP). The final recombinant protein was concentrated to ∼20 mg/mL, flash-frozen in liquid nitrogen, and stored as aliquots at −80 °C for further usage.

#### Micronuclei purification

Micronuclei purification protocol was performed as previously described^8,37,90^. HeLa or 3×FLAG-TREX1 overexpressing HEK293T (both lines expressing H2B-mCherry and GFP-cGAS) cells were treated with 0.5 µM reversine 48 hours prior to harvesting. 10^8^-10^9^ cells were harvested and washed twice in DMEM without serum. Washed cells were resuspended in pre-warmed (37° C) DMEM without serum supplemented with cytochalasin B (Cayman) at 10 µg/mL at a concentration of 10^7^ cells/mL DMEM and incubated at 37° C for 30 minutes. Cells were centrifuged at 300 x g for 5 minutes and cell pellet was resuspended in cold lysis buffer (10 mM Tris-HCl, 2 mM Mg-acetate, 3 mM CaCl_2_, 0.32 M sucrose, 0.1 mM EDTA, 0.1 % (v/v) NP-40, pH 8.5) freshly complemented (with 1 mM dithiothreitol, 0.15 mM spermine, 0.75 mM spermidine, 10 µg/mL cytochalasin B and protease inhibitors) at a concentration of 2 × 10^7^ cells/mL lysis buffer. Resuspended cells were then dounce homogenized by 10 strokes with a loose-fitting pestle. Cell lysates were then mixed with an equal volume of ice cold 1.8 M sucrose buffer (10 mM Tris-HCl, 1.8 M sucrose, 5 mM Mg-acetate, 0.1 mM EDTA, pH 8.0) freshly complemented (with 1 mM dithiothreitol, 0.3 % BSA, 0.15 mM spermine, 0.75 mM spermidine) before use. 10 mL of this mixture (lysed cells + 1.8 M sucrose buffer) was then layered on top of a two-layer sucrose gradient (prepared by slowly adding 20 mL of 1.8 M sucrose buffer slowly on top of 15 mL 1.6 M sucrose buffer in a 50 ml conical tube). This mixture was then centrifuged in a JS-5.2 swinging bucket rotor (Beckman) at 944 × *g* for 20 minutes at 4 °C. Generally, fractions were collected as follows: upper 2 mL contain debris and is discarded; next 5–6 ml contains micronuclei and is collected; final 38 ml contains primary nuclei and is discarded. Fractions containing micronuclei were pooled and diluted 1:5 with FACS buffer (ice cold PBS supplemented with 0.3 % BSA, 0.1 % NP-40 and protease inhibitors). Diluted micronuclei were then centrifuged at 944 × g in JS-5.2 swinging bucket rotor for 20 minutes at 4 °C. Supernatant was then removed by aspiration and micronuclei were resuspended in 2–4 mL of FACS buffer supplemented with 2 µg/mL DAPI. Resuspended samples were filtered through a 40 µm ministrainer (PluriSelect) into FACS tubes. Micronuclei were then sorted by FACSAria (BD Biosciences) into FACS buffer at the MSKCC Flow Cytometry Core Facility. Default FSC and DAPI thresholds were lowered and a log scale was used to visualize micronuclei. Sorted micronuclei were centrifuged at 4000 × *g* in JS-5.2 swinging bucket rotor for 20 minutes at 4 °C and the pellets were stored at -80 °C before lysis for Western blotting or used directly for immunofluorescence microscopy.

#### Proteomic analysis

##### Proteome purification

Unless otherwise specified, all chemicals were from Sigma-Aldrich at the highest available purity. Lanes containing SDS PAGE-resolved proteins were excised from the gel and de-stained using 100 µL of 30 mM potassium hexa-cyanoferrate (III) /100 mM sodium thiosulfate. Destained slabs were washed trice with 200 µL of 25 mM ammonium bicarbonate pH 8.4 / 25 % acetonitrile (v/v, Optima LC/MS grade, Fisher Scientific, Fair Lawn, NJ) and dehydrated using a vacuum centrifuge. Proteins were reduced (10 mM dithiothreitol, 100 mM ammonium bicarbonate pH 8.4, 56 °C for 60 minutes) and alkylated (55 mM iodoacetamide, 100 mM ammonium bicarbonate pH 8.4, 25 °C for 30 minutes, dark), and excess iodoacetamide was removed by three cycles of dehydration (100 µL C_2_H_3_N) and rehydration (100 µL 100 mM ammonium bicarbonate pH 8.4) prior to final dehydration in a vacuum centrifuge. Gel slabs were re-hydrated using 50 mM ammonium bicarbonate pH 8.4 containing 0.04 µg sequencing grade modified porcine trypsin (Promega, Madison, WI) per reaction and proteolysis was allowed to proceed for 16 hours at 37 °C. Tryptic peptide elution from the polyacrylamide slabs was achieved by two consecutive 30 minute incubations in 1 % formic acid, 70 % acetonitrile (v/v) under continuous shaking. Eluates from each sample were pooled, lyophilized and stored at -80 °C until analysis.

##### LC-MS analysis

Peptide pellets were resuspended in 30 µl 0.1% formic acid (99+%, Thermo Scientific, Rockford, IL) and 10% of the solution was analyzed by LC/MS. The LC system consisted in a vented trap-elute setup (Ekspert nanoLC 425, Eksigent, Redwood city, CA) coupled to the Orbitrap Fusion mass spectrometer (Thermo, San Jose, CA) via a nano electro-spray DPV-565 PicoView ion source (New Objective, Woburn, MA). The trap column was fabricated capping a 5 cm × 100 µm internal diameter silica capillary (Polymicro Technologies, Phoenix, AZ) with a 2 mm silicate frit, and pressure loaded with Poros R2-C18 10 µm particles (Life Technologies, Norwalk, CT). The analytical column consisted of a 40 cm × 75 µm internal diameter column with integrated electrospray emitter (New Objective), and was packed with ReproSil-Pur C18-AQ 3 µm particles (Dr. Maisch, Ammerbuch-Entringen, Germany). Samples were loaded on the trap column at 2.5 µl/ minute with the analytical column excluded from the flow path. The analytical column was then put in-line with the trap, and peptides were resolved over 90 minutes using a 5-40% gradient acetonitrile/ 0.1% formic acid (buffer B) gradient in water/0.1% formic acid (buffer A) at 300 nl/minute. The voltage between the mass spectrometer inlet and the electrospray emitter (3 µm internal diameter, fabricated in house from 20 µm internal diameter fused silica capillary), was decreased from 1800 to 1600V in 50V steps at 30, 50, 70, and 90 minutes from contact closure. Precursor ion scans were recorded over the 400–2000 m/z range using the Orbitrap detector (240,000 resolution at m/z 200) with automatic gain control target set 3.2 x 10^5^ ions and maximum injection time of 50 ms. We used data-dependent mass spectral acquisition with monoisotopic precursor selection, precursor selection range 400-2000 Th for ions with charge 2-6, dynamic exclusion (30 sec, 20 ppm tolerance), HCD fragmentation (normalized stepped collision energy 28,32,36 isolation window 1.2 Th) using the top speed algorithm with a duty cycle of 3 sec. Product ion spectra were recorded in the orbitrap trap at 30,000 resolution (automatic gain control= 10^4^ ions, maximum injection time = 54 ms). The mass spectrometry proteomics data have been deposited to the ProteomeXchange Consortium via the PRIDE partner repository with the dataset identifier PXD062583.

#### LC-MS data analysis

Raw files were analyzed using MaxQuant Version 11.6.0.16 and searched against the UniProt human proteome database (as of May 2019, containing isoforms) supplemented with contaminant cRAP database. FDR was set at <0.01 for both peptide and protein level. Mass tolerance was set at 5 ppm for MS1 and 20 ppm for MS2. C-carbamidomethylation was set as fixed modification, while acetylation of protein N-terminus, NQ-deamidation and M-oxidation were allowed as variable modifications. Protein quantification was retrieved from the proteinGroups.txt table, after removing decoy identifications and common contaminants matching the cRAP database (with the exception of trypsin, used as control, and GFP).

**Video S1** Live-cell imaging of HeLa iCas9 sgBAF expressing GFP-TREX1, NLS-RFP, and H2B-iRFP treated with dmso (BAF proficient). 10 minutes interval, scale bar, 10 µm.

**Video S2** Live-cell imaging of HeLa iCas9 sgBAF expressing GFP-TREX1, NLS-RFP, and H2B-iRFP treated with doxycycline (BAF deficient). 10 minutes interval, scale bar, 10 µm.

**Video S3** Live-cell imaging of HeLa iCas9 sgBAF expressing GFP-RPA, NLS-RFP, and H2B-iRFP treated with dmso (BAF proficient). 10 minutes interval, scale bar, 10 µm.

**Video S4** Live-cell imaging of HeLa iCas9 sgBAF expressing GFP-RPA, NLS-RFP, and H2B-iRFP treated with doxycycline (BAF deficient). 10 minutes interval, scale bar, 10 µm.

Supplementary Table S1. Proteomic analysis of purified micronuclei in HeLa cells, related to Figure 1.

Supplementary Table S2. Proteomic analysis of purified micronuclei in HEK293T cells, related to Figure S1.

---

**Figure S1.**
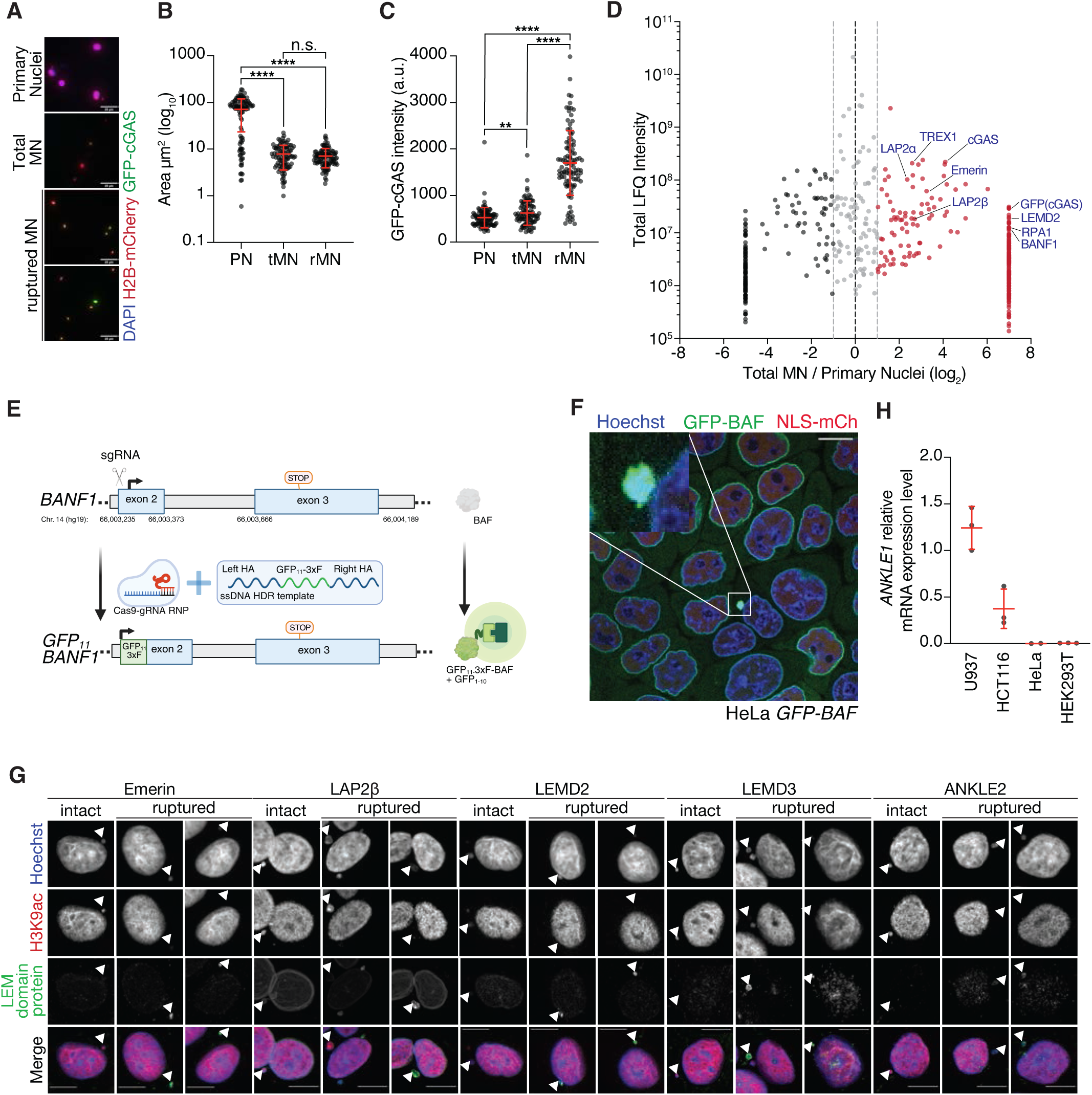
Proteomics of ruptured micronuclei, related to Figure 1. (A) Immunofluorescence for mCherry (H2B) and GFP (cGAS) in the indicated fractions after FACS in HEK293T 3XFlag-TREX1 GFP-cGAS mCherry-H2B. Scale bar, 25 µm. (B) Measurement of the areas of sorted primary nuclei (PN) and micronuclei (MN) sorted using FACS. (C) Quantification of the GFP-cGAS signal intensity with mean and SD as shown in (A). (D) Volcano plots of total LFQ (Label-Free Quantification) intensities plotted against the log₂-transformed ratios of peptide LFQ intensities in ruptured versus total micronuclei in HEK293T 3XFlag-TREX1 H2B-mCherry GFP-cGAS. For proteins specifically detected in either sample, relative abundance was imputed with fixed values (7 and -5 for proteins specifically detected in total micronuclei and primary nuclei, respectively) arbitrarily chosen at the extremes of the empirical ratio distribution. (E) Schematics of targeting GFP_11_ to BAF N terminus in HeLa GFP_1-10_ cells. (F) Live-cell imaging of HeLa GFP_1-10_ GFP_11_-BAF stained with Hoechst. Scale bar, 10 µm. (G) Immunofluorescence of LEM domain proteins in HeLa cells. H3K9Ac serves as a nuclear envelope integrity marker. Scale bar, 10 µm.All *P* values were calculated using one-way ANOVA with Tukey’s multiple-comparisons test (\*\**P* < 0.01; \*\*\*\**P* < 0.0001; n.s., not significant).

**Figure S2.**
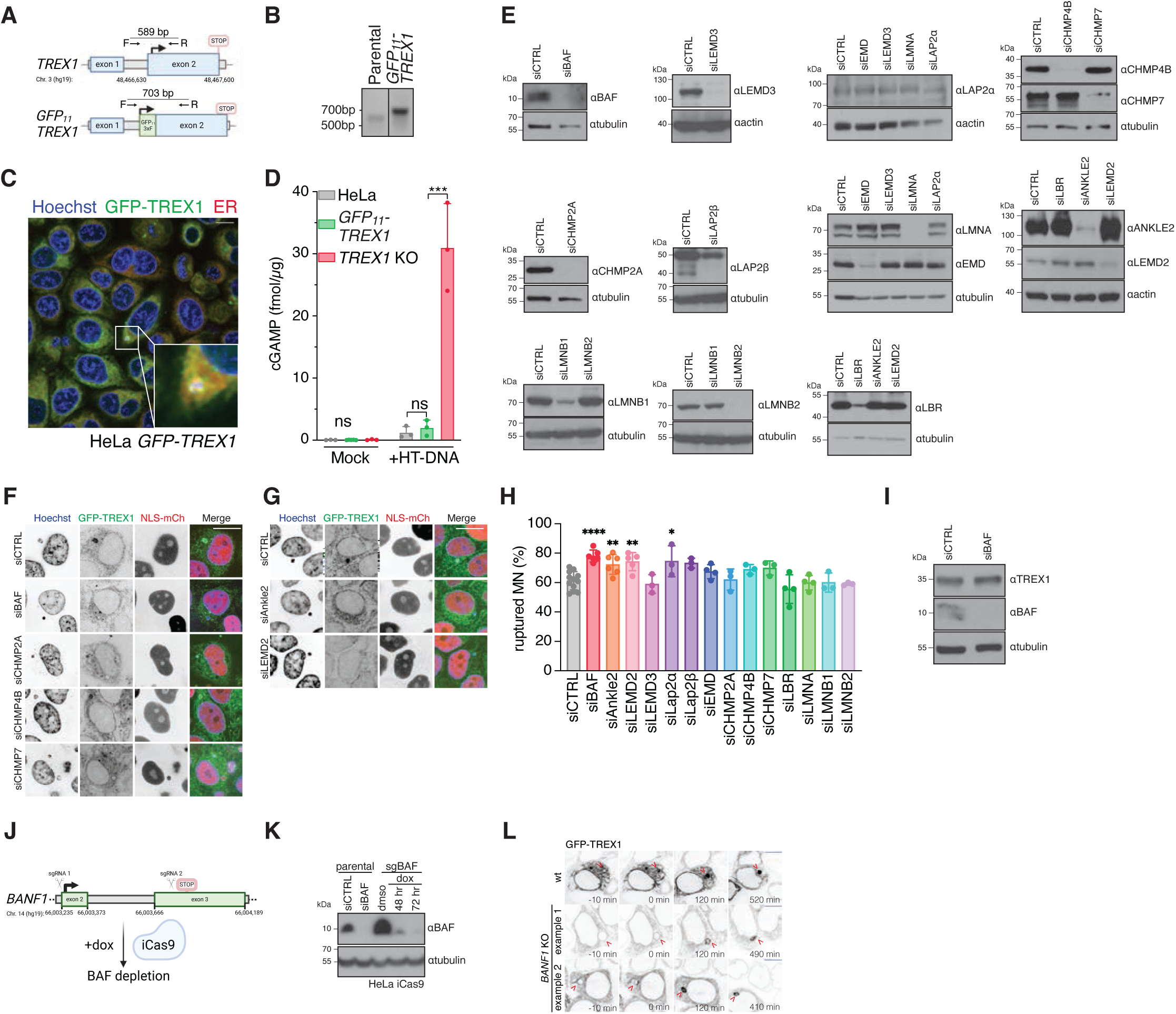
BAF promotes TREX1 recruitment, related to Figure 2. (A) Schematics of PCR validation of GFP_11_ knockin to TREX1 N terminus in HeLa GFP_1-10_ cells. Successful knockin leads to a 703 bp PCR product. (B) PCR validation of HeLa GFP_1-10_ GFP_11_-TREX1 clone compared to parental. (C) Live-cell imaging of HeLa GFP_1-10_ GFP_11_-TREX1 co-stained with ER tracker Red and Hoechst. Scale bar, 10 µm. (D) ELISA analysis of cGAMP production in the indicated cells. Mean ± s.d. of *n* = 3 experiments are shown. Ordinary one-way ANOVA with Dunnett’s multiple comparisons test (ns, not significant; \*\*\**P* < 0.001). (E) Immunoblotting of siRNA screen targets to demonstrate depletion of targets by siRNA. (F) Live-cell imaging of HeLa GFP_1-10_ GFP_11_-TREX1 NLS-RFP upon depletion of BAF or ESCRT-III components by siRNA. Scale bar, 10 µm. (G) Live-cell imaging of HeLa GFP_1-10_ GFP_11_-TREX1 NLS-RFP upon depletion of LEM domain proteins by siRNA. Scale bar, 10 µm. (H) Quantification of micronuclei rupture rate in siRNA screen in Figure 2D. Mean ± s.d.; \*\*\**P* < 0.00=01, \*\**P* < 0.01, \**P* < 0.1,ordinary one-way ANOVA with Dunnett’s multiple comparisons test; *n* = ≥3 independent experiments for each siRNA treatment. (I) Immunoblotting for TREX1 and BAF in HeLa parental cells treated with the indicated siRNA. (J) Schematics of generating inducible BAF KO in HeLa iCas9 cells. Two gRNAs targeting start and end of *BANF1* were stably expressed in HeLa iCas9 cells. Addition of doxycycline leads to Cas9 expression, gRNA cleavage, and BAF depletion. (K) Immunoblotting for BAF in HeLa iCas9 sgBAF cells with the indicated treatment compared to HeLa parental cells treated with the indicated siRNA. (L) Live-cell time-lapse imaging of HeLa inducible *BANF1* KO stably expressing GFP-TREX1 after 72 hours treatment of dmso (*BANF1* WT) or doxycycline (*BANF1* KO), captured at 10-minute intervals, as in Figure 2H. Two examples of *BANF1* KO 7∼8 hours post MN rupture compared to *BANF1* WT. Scale bar, 10 µm.

**Figure S3.**
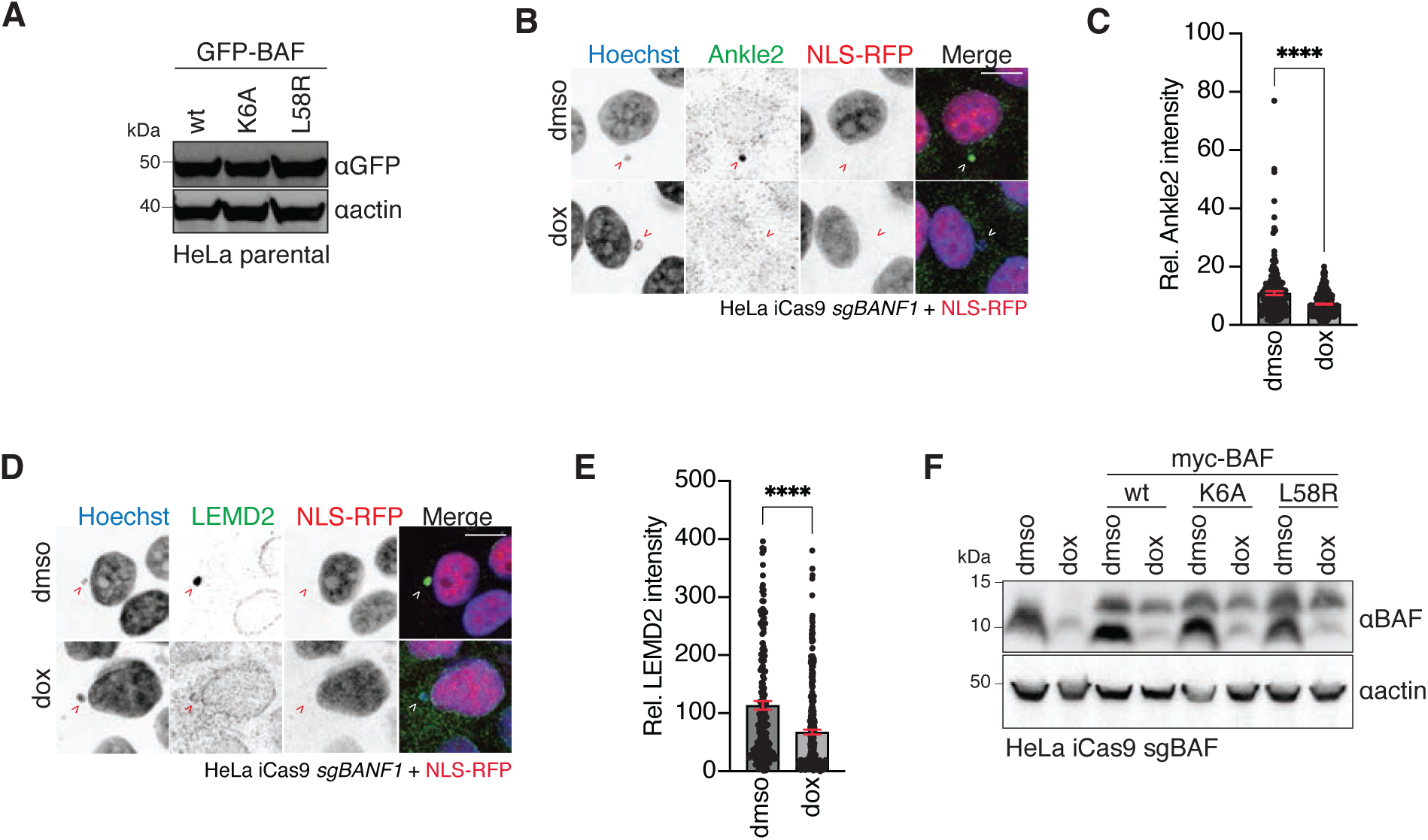
BAF-LEM domain interactions are necessary for TREX1-ER recruitment, related to Figure 3. (A) Immunoblotting for GFP in HeLa parental cells stably expressing the indicated EGFP-BAF-variants. (B) Immunofluorescence of Ankle2 in HeLa iCas9 sgBAF expressing NLS-RFP. Arrows marked ruptured micronuclei indicated by the loss of NLS-RFP signal. Scale bar, 10 µm. (C) Quantification of normalized Ankle2 intensity at ruptured micronuclei as in (B). Mean and SEM of n = 3 independent experiments with greater than 100 total micronuclei quantified per experiment. (D) Immunofluorescence of LEMD2 in HeLa iCas9 sgBAF expressing NLS-RFP. Arrows marked ruptured micronuclei indicated by the loss of NLS-RFP signal. Scale bar, 10 µm. (E) Quantification of normalized LEMD2 intensity at ruptured micronuclei as in (B). Mean and SEM of n = 3 independent experiments with greater than 100 total micronuclei quantified per experiment. (F) Immunoblotting for BAF in HeLa iCas9 sgBAF stably expressing the indicated Myc-BAF-variants and treated with dmso or dox for 72 h. All *P* values were calculated by Student’s *t*-test (****P < 0.0001).

**Figure S4.**
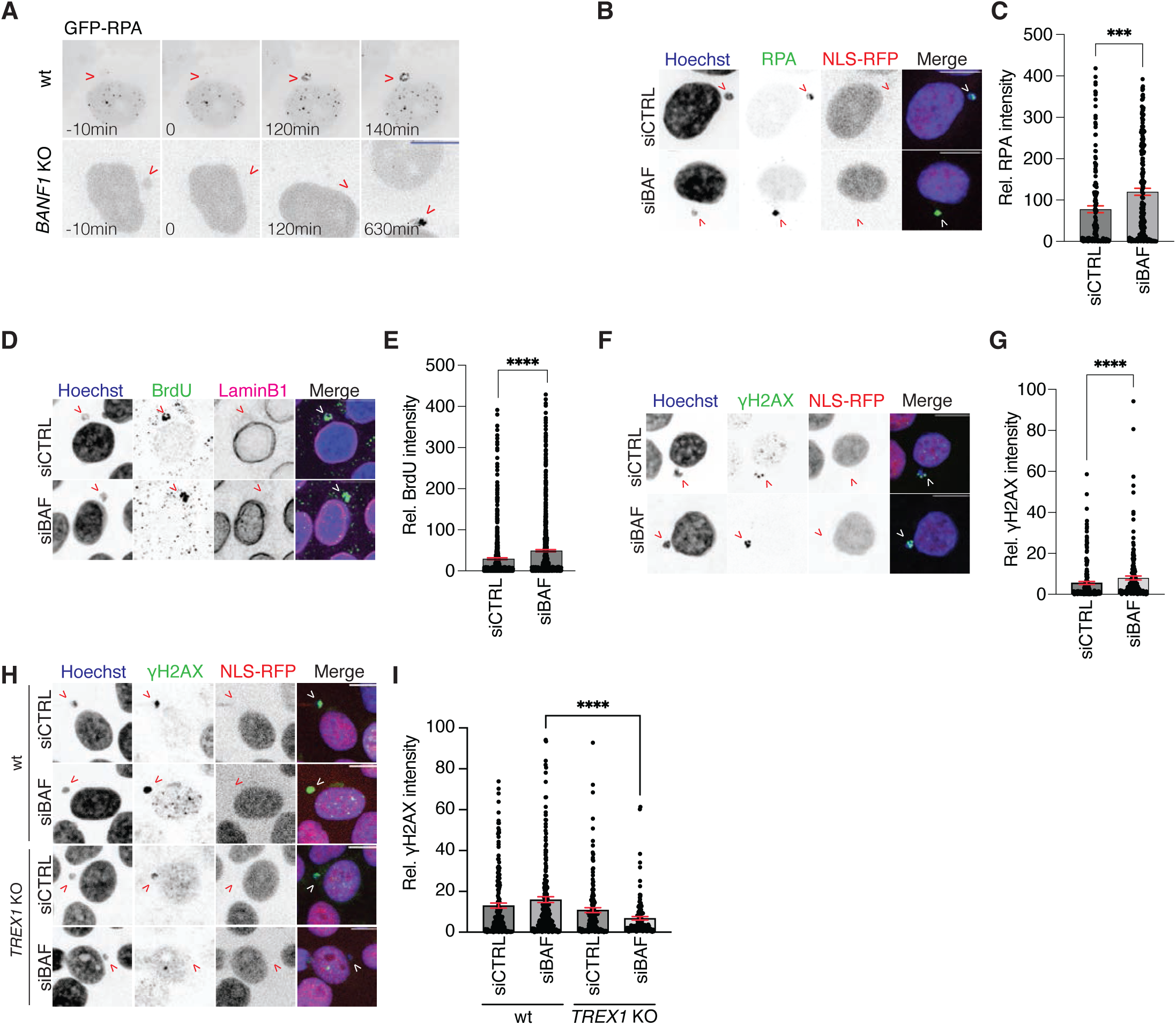
Enhanced TREX1 DNA degradation and ER bypass in BAF-deficient cells, related to Figure 4. (A) Live-cell time-lapse imaging of HeLa inducible BAF KO stably expressing GFP-RPA after 72 hours treatment of dmso (*BANF1* WT) or doxycycline (*BANF1* KO), captured at 10-minute intervals as in Figure 4A. Arrow indicated a micronucleus going through rupture. An example of *BANF1* KO ∼10 hours post MN rupture compared to *BANF1* WT. Scale bar, 10 µm. (B) Immunofluorescence of RPA in HeLa parental cells expressing NLS-RFP treated with CTRL or BAF siRNA. Arrows marked ruptured micronuclei indicated by the loss of NLS-RFP signal. Scale bar, 10 µm. (C) Quantification of normalized RPA intensity in ruptured micronuclei as in (B). (D) Immunofluorescence of native BrdU in HeLa parental cells expressing NLS-RFP treated with CTRL or BAF siRNA. Arrows marked ruptured micronuclei indicated by the loss of LaminB1 signal. Scale bar, 10 µm. (E) Quantification of normalized BrdU intensity in ruptured micronuclei as in (D). (F) Immunofluorescence of γH2AX in HeLa parental cells expressing NLS-RFP treated with CTRL or BAF. Arrows marked ruptured micronuclei indicated by the loss of NLS-RFP signal. Scale bar, 10 µm. (G) Quantification of normalized γH2AX intensity in ruptured micronuclei as in (F). (H) Immunofluorescence of γH2AX in HeLa TREX1 KO treated with CTRL or BAF siRNA. Arrows marked ruptured micronuclei indicated by the loss of NLS-RFP signal. Scale bar, 10 µm. (I) Quantification of normalized γH2AX intensity in ruptured micronuclei as in (H). All *P* values were calculated by Student’s *t*-test (***P < 0.001, ****P < 0.0001). Mean ± s.e.m. of *n* = 3 independent experiments with greater than 100 total micronuclei quantified per experiment.

**Figure S5.**
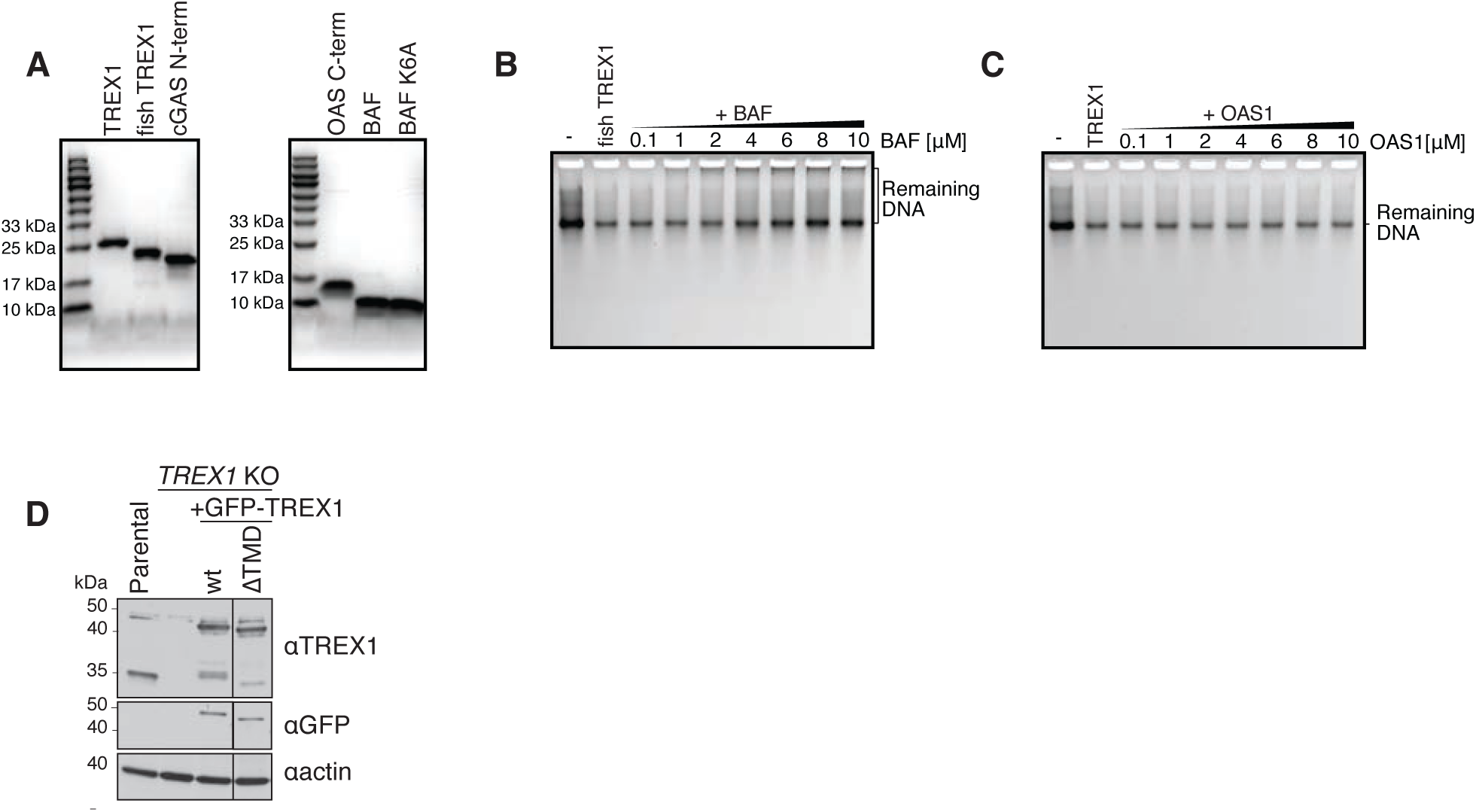
BAF regulates TREX1 DNA degradation *in vitro*, related to Figure 5. (A) SDS-page gel of purified proteins used in *in vitro* nuclease assays shown in Figures 5A-C. (B) Representative DNA gel from *in vitro* nuclease assay. A dsDNA substrate was co-incubated with a fixed concentration of purified fish TREX1 (*Labeo rohita)* protein together with the indicated concentration of purified human BAF protein. - = no TREX1 added. (C) Representative DNA gel from *in vitro* nuclease assay. A dsDNA substrate was co-incubated with a fixed concentration of purified human TREX1 protein together with the indicated concentration of purified human OAS1 C-terminus protein. - = no TREX1 added. (D) Immunoblotting of HeLa parental and TREX1 KO complemented with GFP-TREX1 FL or ΔTMD as in Figure 5F. Note that FL or ΔTMD transgene expression experiences TREX1 clipping that is absent in *TREX1* KO.

**Figure S6.**
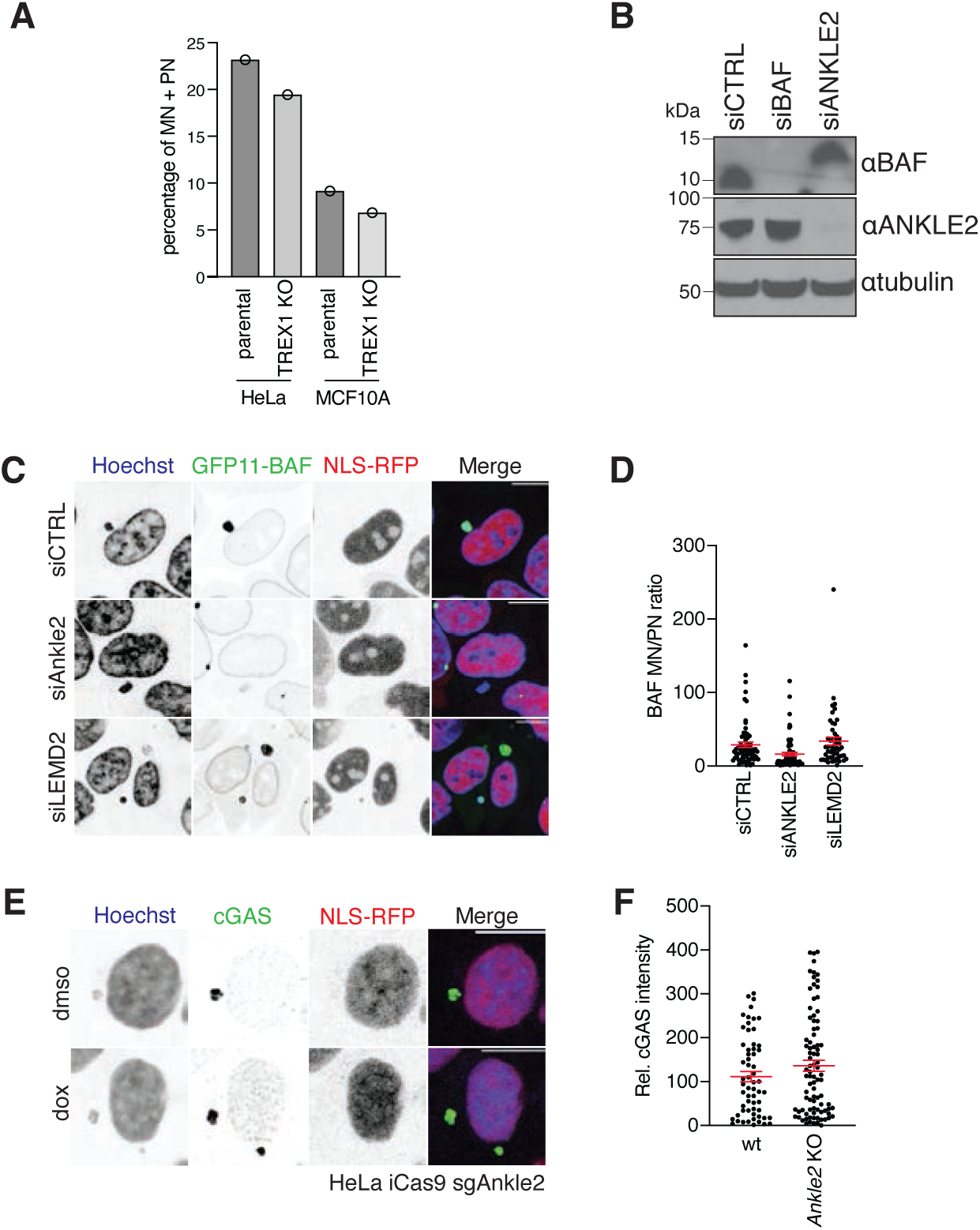
BAF inhibits cGAS activation at ruptured micronuclei, related to Figure 6. (A) Immunoblotting for BAF and Ankle2 in HeLa parental cells treated with the indicated siRNA. (B) Live-cell imaging of HeLa GFP_1-10_ GFP_11_-BAF expressing NLS-RFP treated with the indicated siRNA. Scale bar, 10 µm. (C) Quantification of normalized fluorescence signal intensities of GFP-BAF MN/PN ratio. Mean ± s.e.m *n* = 1. More than 100 total micronuclei were analyzed. (D) Immunofluorescence of cGAS in HeLa iCas9 sgAnkle2 cells expressing NLS-RFP treated with dmso or dox for 72 hours. Scale bar, 10 µm. (E) Quantification of normalized cGAS intensity in ruptured micronuclei as (D). Mean ± s.e.m *n* = 1. More than 100 total micronuclei were analyzed. (F) Quantification of micronucleation rate in HeLa and MCF10A in the indicated genotypes. One independent experiment with more than 1000 cells analyzed.

